# Nutraceuticals Silybin B, resveratrol and epigallocatechin-3 gallate (EGCG) bind to cardiac muscle troponin to restore the loss of lusitropy caused by cardiomyopathy mutations *in vitro, in vivo*, and *in silico*

**DOI:** 10.1101/2024.05.09.593307

**Authors:** Zeyu Yang, Alice Sheehan, Andrew Messer, Sharmane Tsui, Alexander Sparrow, Charles Redwood, Vladimir Kren, Ian R. Gould, Steven B. Marston

**Affiliations:** Institute of Chemical Biology, Molecular Sciences Research Hub and Department of Chemistry, Molecular Sciences Research Hub, Imperial College London, London W12 0BZ, U.K.; NHLI, Imperial College London, London, W12 ONN, United Kingdom; Division of Cardiovascular Medicine, Radcliffe Department of Medicine, University of Oxford, Oxford, United Kingdom and British Heart Foundation Centre of Research Excellence, University of Oxford, Oxford, United Kingdom; https://orcid.org/0000-0002-3354-3314; Division of Cardiovascular Medicine, Radcliffe Department of Medicine, University of Oxford, Oxford, United Kingdom and British Heart Foundation Centre of Research Excellence, University of Oxford, Oxford, United Kingdom; https://orcid.org/0000-0002-8673-7542; Laboratory of Biotransformation, Institute of Microbiology of the Czech Academy of Sciences, Prague, Czechia https://orcid.org/0000-0002-1091-4020.; Department of Chemistry, Molecular Sciences Research Hub and Institute of Chemical Biology, Molecular Sciences Research Hub, Imperial College London, London W12 0BZ, U.K.; https://orcid.org/0000-0003-3559-0234; National Heart & Lung Institute, Imperial College London, London W12 0NN, U.K. https://orcid.org/0000-0001-6054-6070

## Abstract

Adrenergic activation of protein kinase A (PKA) targets the thin filaments of the cardiac muscle, specifically phosphorylating cTroponin I Ser22 and Ser23, causing a higher rate of Ca^2+^ dissociation from cTnC leading to a faster relaxation rate (lusitropy). This modulation is often suppressed by mutations that cause cardiomyopathy (uncoupling) and this could be sufficient to induce cardiomyopathy. A drug that could restore the phosphorylation-dependent modulation of relaxation rate could have the potential for treatment of these pathologies.

We found, using single thin filament *in vitro* motility assays that the small molecules including silybin B, resveratrol, and epigallocatechin-3 gallate (EGCG) can restore coupling.

We performed molecular dynamics simulations of the unphosphorylated and phosphorylated cardiac Troponin core with the TNNC1 G159D mutation. We found that silybin B, EGCG, and resveratrol restored the phosphorylation-induced change in the TnC helix A/B angle and the interdomain angle to wild-type values, whilst silybin A and epicatechin gallate (ECG) did not. In unphosphorylated G159D the recoupling molecules were observed to be frequently intercalated between The N terminal peptide of Troponin I and troponin C. In contrast, the controls, silybin A, and ECG bound to the surface. All of the interactions were diminished when troponin I was phosphorylated.

We also performed studies with intact transgenic ACTC E99K mouse cells and TNNT2 R92Q-transfected guinea pig cardiomyocytes. The mutations blunt the increase in relaxation speed due to dobutamine; resveratrol, EGCG, and silybin B could restore the dobutamine response whilst silybin A did not. Thus recoupling by small molecules is demonstrated *in vitro, in vivo*, and *in silico*.

## INTRODUCTION

In cardiac muscle, contractility is controlled by changes in the concentration of intracellular Ca^2+^ ion that binds to troponin, a component of the thin filaments, to activate the contractile interaction of thick and thin filaments. In addition to this Ca^2+^ switch, cardiac muscle possesses a unique modulatory mechanism that allows the heart to meet increased oxygen demand during exercise.

Release of adrenaline and noradrenaline activates *β*-1 receptors in cardiac myocytes and leads to activation of adenylyl cyclase *via* stimulatory G-protein (Gs). The resulting increase of the cytosolic cyclic adenosine monophosphate (cAMP) levels leads to activation of protein kinase A (PKA), which phosphorylates a number of targets in the sarcolemma, sarcoplasmic reticulum, and contractile apparatus to increase contractile force and heart rate. In the thin filaments of the contractile apparatus, troponin I (cTnI) is the target for PKA phosphorylation and is phosphorylated at residues Ser22 and Ser23 in the cardiac-specific N-terminal peptide (NcTnI: residues 1 to 32). Phosphorylation is *coupled* to a 2-3 fold decrease of affinity of cTn for Ca^2+^ due to altered cTnC-cTnI interactions, linked to a higher rate of Ca^2+^ dissociation from cTnC (1-4). The consequent faster relaxation rate of the cardiac muscle due to TnI phosphorylation (lusitropy) is essential for shortening the cardiac muscle contraction-relaxation cycle allowing for efficient contraction at a faster heart rate.

Many *in vitro* studies have demonstrated that mutations in thin filament proteins associated with inherited cardiomyopathies (hypertrophic cardiomyopathy, HCM, and dilated cardiomyopathy, DCM) abolish the relationship between Ca^2+^ sensitivity and TnI phosphorylation by PKA and that the mutations also impair lusitropy in many animal models (5). This trait, which we have termed ‘*uncoupling*’, was first noted in 2001 (6,7). In 2007-8 three publications studying the TNNC1 G159D mutation (G159D), associated with DCM, showed uncoupling with recombinant mutant troponin, with troponin extracted from a patient with the mutation and with rat trabeculae with the mutation exchanged in myocytes (8-10). Subsequently, almost every thin filament mutation that was tested proved to be uncoupled including the HCM mutations TPM1 E180G, TNNT2 R92Q, and ACTC E99K (HCM) used in this study (summarized in (5,11)). It is relevant to note that in *in vitro* experiments, uncoupling is always complete, irrespective of the mutation and that uncoupling is associated with both HCM and DCM-causing mutations. Uncoupling results in blunting of the lusitropic response to β-1 receptor stimulation. Suppression of lusitropy is linked to heart failure phenotypes in animal models and, clinically, reduced response to adrenergic stimulation is correlated with cardiac adverse outcome. (5,12,13).

A drug that could restore the phosphorylation-dependent modulation of Ca^2+^- sensitivity could have potential for treatment of these pathologies. We have investigated small molecules that can specifically reverse these abnormalities *in vitro (‘recoupling’)*. Based on our lead compound, epigallocatechin-3-gallate (EGCG) (14,15) that is both a desensitiser and a recoupler, we examined 40 compounds and found 23 compounds that reversed the uncoupling. Many of these are pure recouplers (i.e. have no desensitising activity) whilst 3 of the compounds desensitized but did not recouple and the rest had no effect (16). Thus recoupling and desensitisation are independent processes. Importantly, recoupling was complete and independent of the causative mutation and the nature of the compound (documented in supplementary material 1). Thus, small molecules that can act as desensitisers or recouplers have the potential to treatments for HCM or DCM. However we do not know the molecular mechanism of the recoupling phenomenon, nor do we know whether the recoupling compounds we investigated *in vitro* are effective in intact muscle. In this paper, we aim to answer these questions *in vitro, in silico, and in vivo*.

Understanding how phosphorylation modulates troponin function is not straightforward since troponin contains several disordered regions that are crucial to regulation and are not accessible to methodologies such as X-ray diffraction or cryoelectron microscopy that only resolve static structures. We recently used molecular dynamics (MD) simulation to determine the atomistic changes that occur when Ser22, 23 of cTnI is phosphorylated in the regulatory core of troponin. These studies showed that phosphorylation induced no significant stable structural changes but rather phosphorylation changed the dynamics of the transitions between a small number of states (17,18). We established metrics characteristic of the Ser22/23 phosphorylation transition: changes in the distribution of the angle between helices A and B of troponin C and the angle between the NcTnC domain and the ITC domain that were consistent with phosphorylation destabilising the open state of TnC. Importantly, the introduction of the TNNC1 G159D mutation profoundly changed these metrics. Unphosphorylated G159D tended to have values equivalent to phosphorylated wild-type and upon phosphorylation the values for G159D moved in the opposite direction to wild-type (18).

The analysis of MD simulations of G159D troponin clearly shows that the mutation abrogates the effects of phosphorylation on structure and dynamics and thus explains its uncoupling activity. We may also use these metrics to determine the effects of small molecules on the system and to test whether recoupling *in vitro* corresponds to the restoration of wild-type dynamics *in silico*.

We analysed molecular dynamics simulations of unphosphorylated and phosphorylated cardiac Troponin core with the G159D mutation in the presence of two pure recouplers, silybin B (SB) and resveratrol, and epigallocatechin-3 gallate (EGCG), a mixed recoupler and desensitiser. These were compared with silybin A (SA), the stereoisomer of silybin B and epicatechin gallate (ECG), closely related to EGCG, that have no measurable recoupling activity *in vitro* as controls. We found that silybin B, EGCG, and resveratrol restored most key metrics towards wild-type values, whilst silybin A (SA) and ECG did not. EGCG gave a complex response consistent with its dual function as recoupler and desensitiser.

In parallel, we have extended our studies to intact cardiomyocytes. Both HCM and DCM-causing mutations that uncouple *in vitro* blunt the response to β1 adrenergic stimulation in intact cardiomyocytes and cardiac tissue (5). We have therefore tested the effects of small molecules, that recouple *in vitro,* on the response to a β1 adrenergic agonist in myocytes with ACTC E99K and TNNT2 R92Q mutations. We found that resveratrol, EGCG, and silybin B could restore the β1 adrenergic response whilst silybin A did not; in the case of EGCG, the response was partly confounded by off-target effects.

## 2. RESULTS

### 2.1 In vitro effects of small molecules on in vitro motility

In previous studies, we used an *in vitro* motility assay to demonstrate that small molecules, including EGCG, silybin B, and resveratrol were able to restore the lack of response to troponin I phosphorylation caused by mutations in thin filament proteins, whereas silybin A and ECG did not (5,19). Figure 1A summarises the effects of these compounds on wild-type thin filaments and upon thin filaments containing the uncoupling HCM mutation TPM1 E180G measured at a single Ca^2+^ concentration (See Supplementary Information 3) in phosphorylated and unphosphorylated states. Figure 1B shows the effects of EGCG, silybin B, and silybin A on the Ca^2+-^regulation of thin filaments containing the TNNC1 (cardiac TnC) G159D mutation. None of the compounds affect the phosphorylation-dependent Ca^2+^-sensitivity shift in wild-type thin filaments. The Ca^2+^ -sensitivity of mutant thin filaments is not modulated by phosphorylation (uncoupled), however, the coupling is restored to both TPM1 E180G and TNNC G159D by 100µM EGCG, resveratrol or silybin B but not by silybin A or ECG (Supplementary Information 2C). In addition, EGCG, ECG, and silybin A are desensitisers, reducing Ca^2+^-sensitivity in wild-type and mutant by about 1.5-fold independently of the recoupling property, thus silybin B and resveratrol are ‘pure’ recouplers whilst EGCG is a mixed desensitiser and recoupler. These results are in accord with the results of previous studies showing that the recoupling effect of small molecules, silybin B, EGCG, and resveratrol is independent of the causative mutation and that a full restoration of the effects of troponin I phosphorylation is achieved by all recouplers (15,16).

### 2.2 Contractility of isolated cardiomyocytes

In order to assess the effects of EGCG, silybin A, and silybin B on uncoupling in intact cells, we isolated cardiomyocytes from wild-type and two well-characterised models of hypertrophic cardiomyopathy that are uncoupled *in vitro* due to mutations in a thin filament protein. The models were ACTC E99K transgenic mouse myocytes and guinea-pig cardiomyocytes with the TNNT2 (TnT) R92Q mutation replacing the wild-type protein, introduced by transfection (19-21).

The contractility of wild-type and mutant myocytes was measured. We studied three main parameters that are modulated by β1 adrenergic activation: the contractile amplitude (% peak height), the time of contraction (time to 90% peak, ttp_90_), and the time of relaxation (time to 90% baseline, ttb_90_), see Figure 2A and Supplementary Information 5. Whilst the contraction amplitude of non-transgenic mouse and guinea-pig myocytes was similar, ttp_90_ and ttb_90_ in guinea pig were about double that of mouse, indicating slower kinetics.

**Figure 2A.**
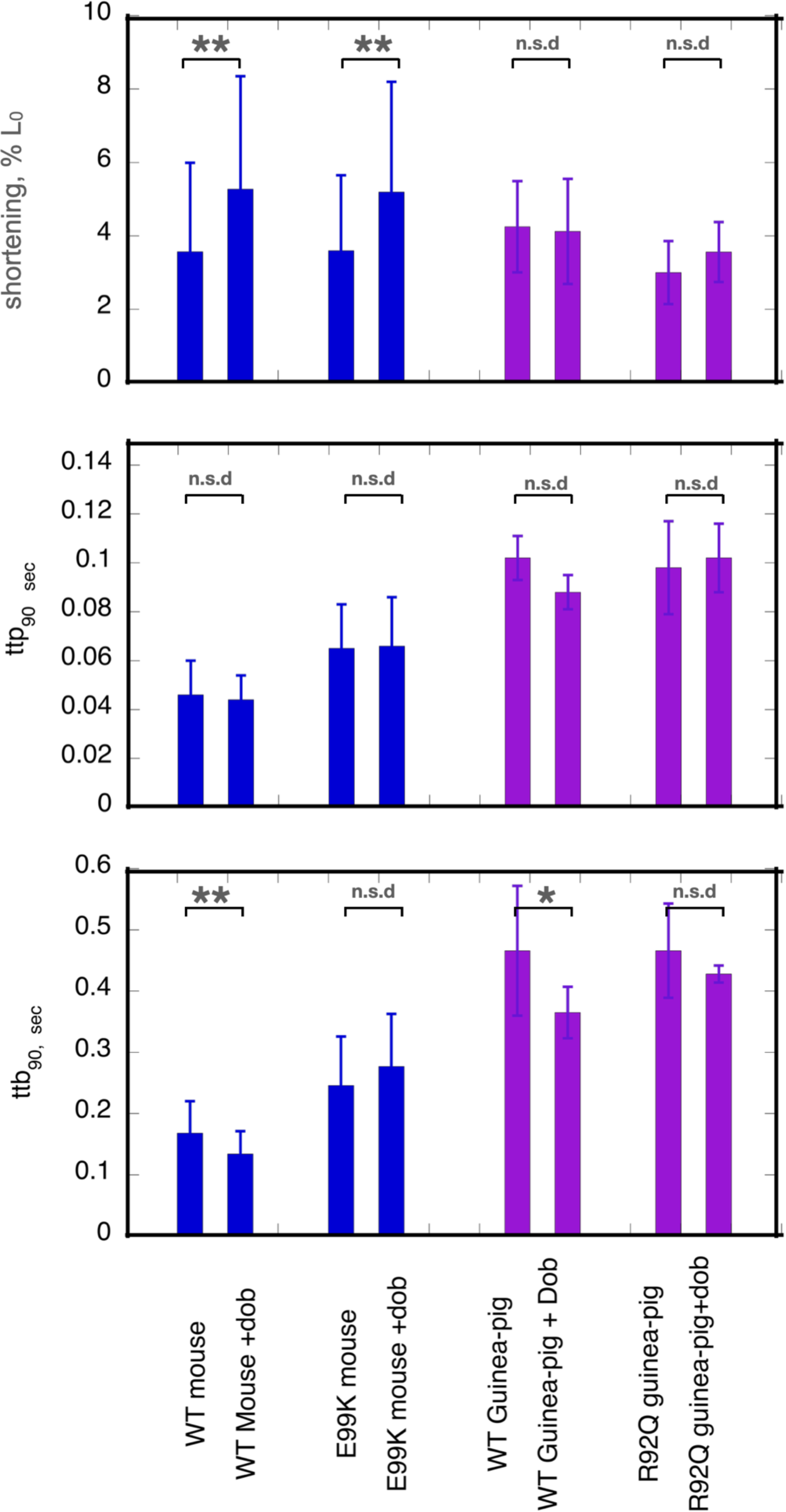
Effects of dobutamine on key parameters of cardiac myocyte contractility. Shortening amplitude, ttp_90_ and ttb_90_ parameters are shown before and after addition of dobutamine. Blue-WT and E99K mouse compared. Purple-WT and R92Q guinea-pig compared. Full data and statistical analysis in supplementary information 5

Cardiomyopathic mutations affected the baseline contractility of cardiomyocytes as previously noted (20,21). Both ttp_90_ and ttb_90_ were greater in ACTC E99K mouse cardiac cells compared to WT cells by 41% and 46% respectively, indicating slower contraction and relaxation whilst contractile amplitude was not different. In the Guinea-pig TNNT2 R92Q cells, ttp_90_ was the same but ttb_90_ was 35% greater than the wild type indicating slower relaxation due to the mutation.

The normal response to β1 receptor activation is increased amplitude of shortening and faster contraction and relaxation (lusitropy). In contrast, mutations that are observed to be uncoupled *in vitro* have a blunted response to β1 adrenergic stimulation in intact cardiomyocytes and cardiac tissue (5). We confirmed this finding in our preparations (Figure 2A).

We treated cells with the β1-selective agonist dobutamine in the presence of the highly selective β2 receptor antagonist, ICI-118,551. In NTG mouse cells, adding dobutamine increased the amplitude of shortening by 47% and decreased ttb_90_ by 21%, thus indicating both inotropic and lusitropic effects. In guinea-pig cells, dobutamine did not affect amplitude but ttb_90_ was decreased by 21% indicating normal lusitropy.

In the mutant cells, we did not observe any significant lusitropy, indicating that β1 adrenergic stimulation and the consequent troponin I phosphorylation were uncoupled from its effect on the rate of relaxation (Supplementary Information 5, Figure 2A). We then tested the ability of small molecules to restore ttb_90_ to WT levels in the presence of dobutamine.

EGCG, silybin A, silybin B, and resveratrol were examined as potential re-coupling compounds. Mouse E99K or guinea-pig R92Q myocytes were incubated in the presence of the small molecules and contractility was measured before and after adding 0.4µM dobutamine. Preliminary titrations indicated that maximal effects of these molecules were achieved with micromolar concentrations, much lower than the concentrations needed for the *in vitro* motility assays. We quantified lusitropy as the fractional change of ttb_90_ due to dobutamine [ 1-(ttb_90_ +Dob/ttb_90_ -dob)]. The baseline lusitropy in NTG mouse or guinea pig was a 20-24% *decrease* whilst in the mutant myocytes dobutamine induced a non-significant *increase* in ttb_90_ (12% in mouse, 9% in guinea pig) (Figure 2A).

In the presence of the known recoupling molecules, there was a clear restoration of the lusitropic effect (Figure 2B). In the presence of silybin B, dobutamine reduced ttb_90_ by 25% in E99K mouse and 13% in R92Q guinea pig. In the presence of resveratrol, dobutamine reduced ttb_90_ by 41% in E99K mouse and 17% in R92Q guinea pig. With EGCG, dobutamine reduced ttb_90_ by 22% in E99K mouse and reduced ttb_90_ by 17% in R92Q guinea pig. In contrast, silybin A, which does not recouple *in vitro*, did not restore lusitropy; we observed a non-significant 4% *increase* in ttb_90_ in E99K mouse and an 18% *increase* in R92Q guinea pig myocytes which is similar to the effect of dobutamine in the absence of small molecules (9% increase), see Figure 2B and Supplementary Information 6. ECG was not tested in this system. Thus, the small molecules that were shown to be recouplers *in vitro* also restored the blunted lusitropic response to β1 adrenergic stimulation in intact mutant myocytes. The specificity of this effect is confirmed by the inability of silybin A to restore lusitropy.

**Figure 2B.**
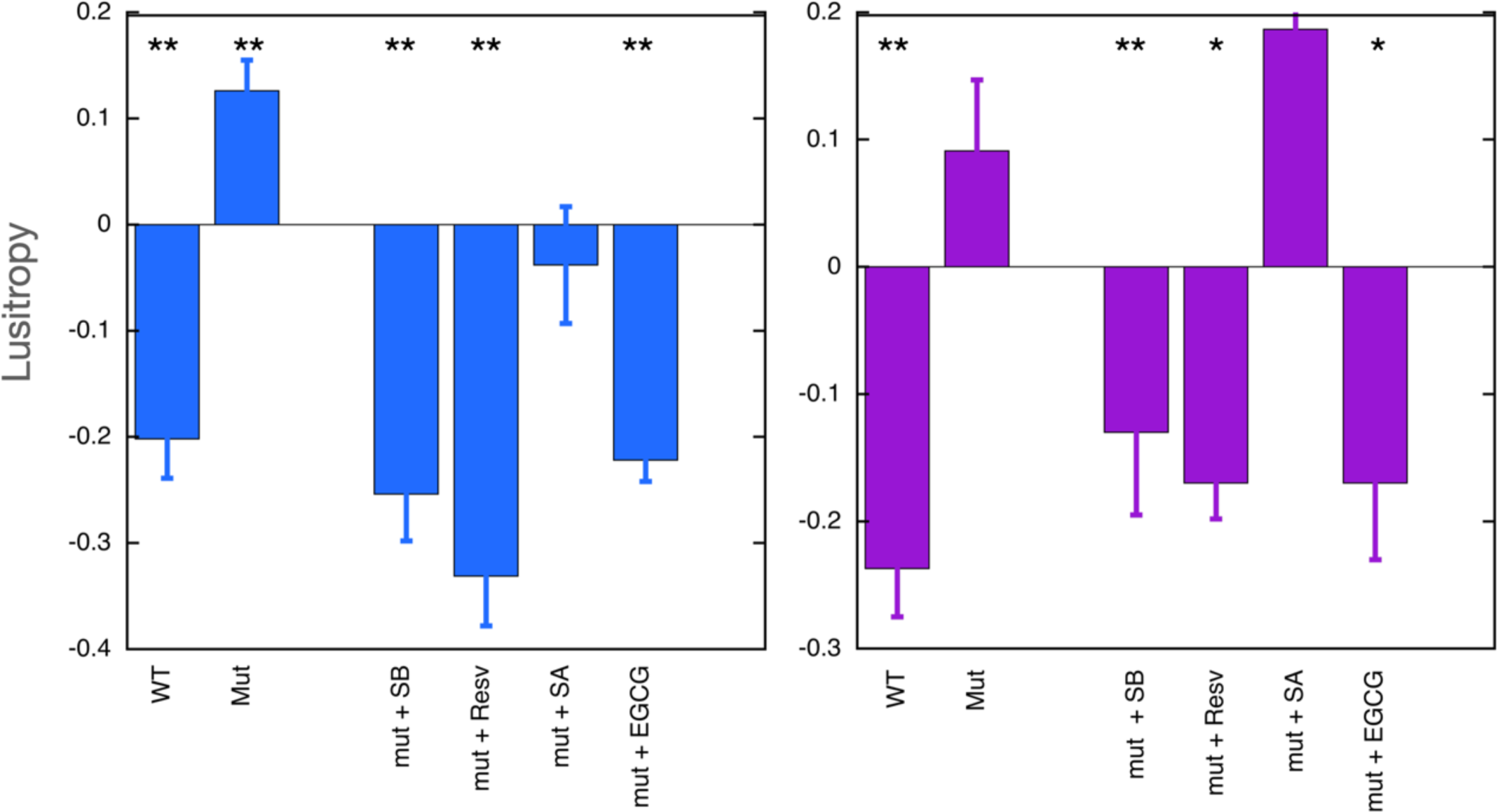
Lusitropy and the effect of small molecules measured in cardiomyocytes. Lusitropy is defined as the fractional change of ttb_90_ due to dobutamine [ 1-( ttb_90_ +Dob/ ttb_90_ -Dob)]. Positive lusitropy (increase in relaxation speed) corresponds to a decrease in this parameter. Blue, mouse experiments, Purple, Guinea-pig experiments. Lusitropy is lost in the mutant myocytes but is restored by EGCG, SilybinB and resveratrol but not by SilybinA. Original data and statistical analysis in supplementary information 6

### 2.3 Effects of small molecules on troponin phosphorylation determined by Molecular Dynamics Simulations

To understand how the small molecules modulated troponin behaviour at the atomic level we performed molecular dynamics simulations (5×1500ns) on phosphorylated and unphosphorylated wild-type and TnC G159D DCM mutant troponin in the presence of silybin A, silybin B, EGCG, ECG, and resveratrol.

#### 2.3.1 Recoupling ligands can restore the phosphorylation-dependent changes in molecular dynamics metrics that are suppressed in TnC G159D mutant

In previous molecular dynamics simulations, we demonstrated that phosphorylation of troponin I does not significantly alter the structure of the troponin core. However, phosphorylation does change the dynamics of troponin and the uncoupling mutant TnC G159D. We have derived several metrics that are characteristic of the phosphorylated and unphosphorylated state. In particular, the distribution of the angle between cTnC helices A and B and the angle between quasi-rigid troponin domains (NcTnC and the ITC domain) may be used since the effect of TnI phosphorylation on the metrics of the uncoupling TnC G159D mutation are strikingly different from wild-type with phosphorylation-dependent changes in G159D being mostly in the opposite direction to those in wild-type [24]. Figure 3A shows the angle distributions comparing WT and G159D troponin whilst Figure 3B shows the effect of small molecules and phosphorylation on the angle distributions. Corresponding data for ECG is in Supplementary Information 2

**Figure 3.**
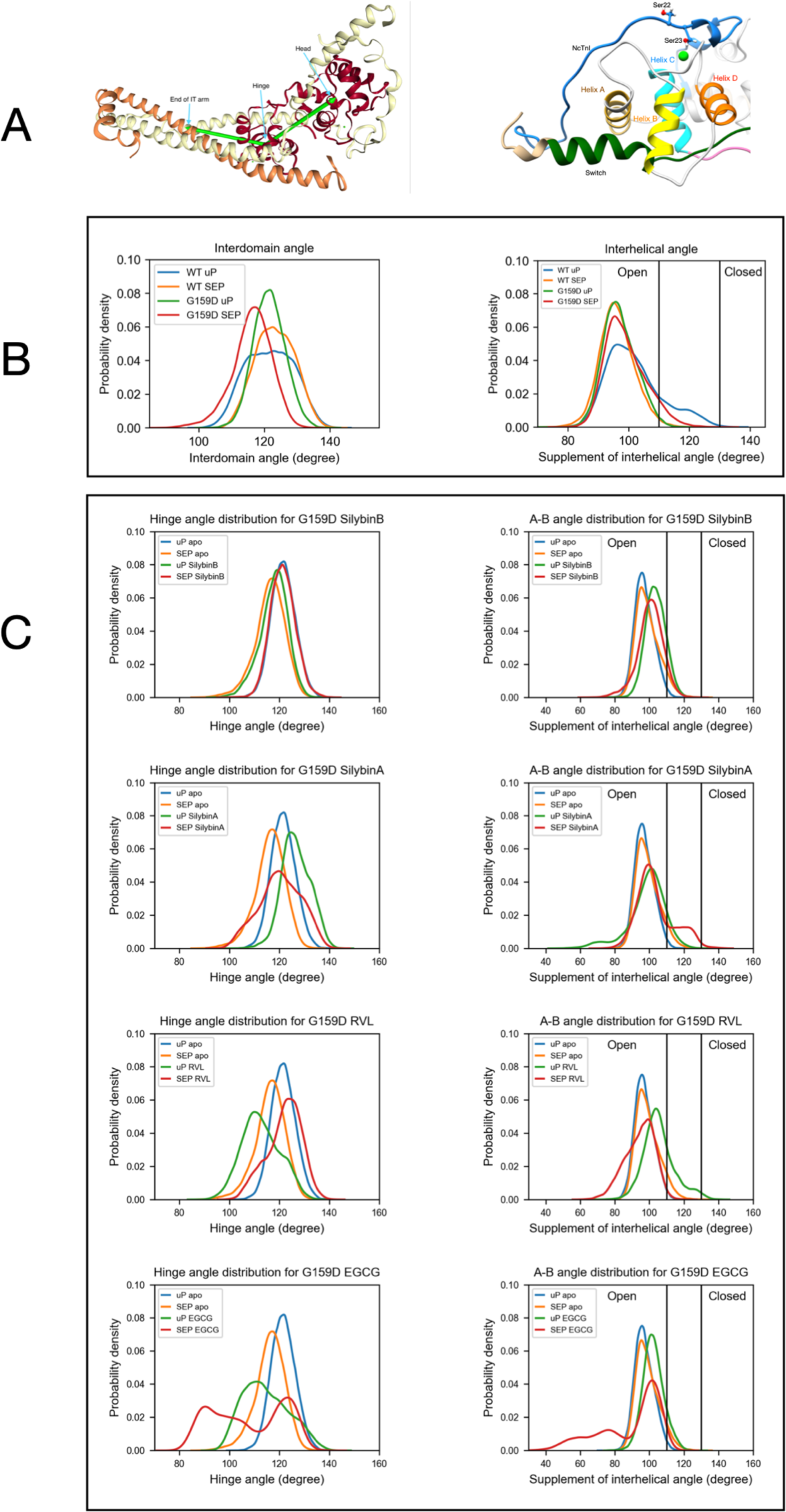
The distribution of a helix A/B and Interdomain angles determined by molecular dynamics simulations. A Left: Model definining the location of the hinge between N-terminal and C-terminal domains of troponin C. Right: Model showing the orientation of helices A and B in the N-terminus of troponin C B The effects of the G159D mutation and phosphorylation on the distribution of the interdomain angle and the heix A/B angle. Unohosphorylated, uP, phosphorylated, SEP. C The effects of phosphorylation and small molecules on the distribution of the interdomain angle (left) and the heix A/B angle (right) For G159D. Table 1 shows the quantification of the distributions. Distribution plots for ECG are shown in supplementary information 2.

#### 2.3.2 Angle between troponin C helices A and B

The A/B helix angle may be used as a metric for the opening and closing of the hydrophobic patch (Table 1A). In WT, the effect of phosphorylation is a decrease of the mean A/B angle, from 101.8° to 96.1° (Δ= - 5.6°). For the G159D mutant we observe the complete inverse behaviour: upon phosphorylation, the A/B angle increases from 97.1° to 99.0° (Δ= +1.9°). Comparing the WT and G159D, unphosphorylated A/B angles we see that the WT angle is significantly higher than G159D.

**TABLE 1A.**
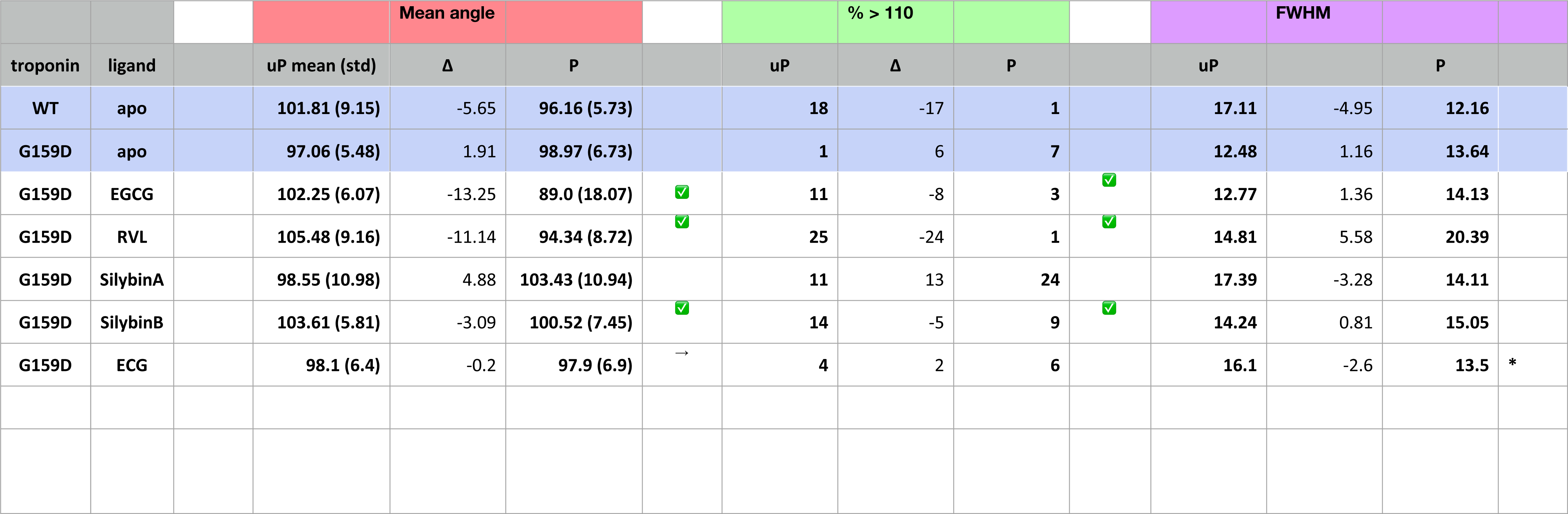
A/B helix angle metrics.

We determined the effect of the small molecule ligands on the interhelix angle distribution in G159D and wild-type troponin (Figure 3, Table 1B). In wild-type the ligands tended to have no effect or to increase A/B angle in both unphosphorylated and phosphorylated states, however, the decrease in mean angle upon phosphorylation, associated with a normal response to phosphorylation was preserved. In G159D the effects of the small molecules were different, the three recouplers restored the decrease in A/B angle on phosphorylation (SB, -3.1°, EGCG, -13.3°, RVL, -11.1° compared with -5.6° in the wild-type) whereas with silybin A, which has no recoupling activity *in vitro,* the change resembled that of G159D ( SA, +4.88°, ECG, -0.2° compared with ApoG159D +1.9°). This pattern of results was also clear when considering the percentage of the area under the curve for the A/B angle above 110°, corresponding to the boundary of the open state of troponin C. With G159D this parameter increased from 1 to 7% on phosphorylation. It increased from 11 to 24% in the presence of silybin A and from 4 to 6% with ECG but with the three recoupling molecules the change was in the opposite direction (SB, 14>9%, EGCG, 11>3%, RVL 25>1% compared with 18 > 1% in wild-type troponin). Thus the three small molecules that are physiological recouplers restore the phosphorylation-dependent changes in the disease-causing, uncoupled, G159D mutation at the atomic level and silybin A and ECG that are not recouplers, do not restore the A/B angle change.

**TABLE 1B.**
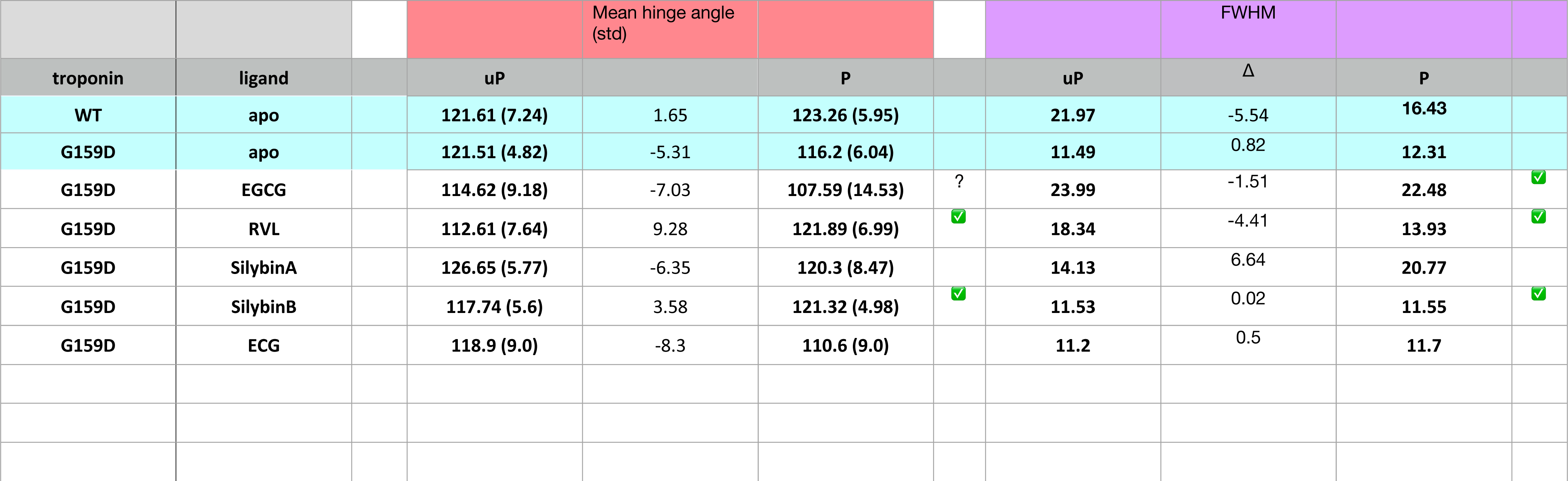
Hinge Angle metrics.

#### 2.3.3 Effect of small molecules on hinge motion

The principal motion of troponin is a hinge-like motion between the two quasi-rigid domains of troponin, NcTnC, and the ITC domain. The distribution of hinge angles in wild-type troponin shows an increase in the mean angle upon phosphorylation and a lengthening of the TnC peptide link between the domains. In contrast, phosphorylation of G159D shifts the inter-domain angle distribution towards a lower mean angle. Thus the wild-type response to phosphorylation is disrupted by the G159D mutation at the atomic level (Figure 3C, Table 1B, Table 2A).

**TABLE 2A.** Interdomain distance metrics.

| Interdomain distance metrics |  |  |  |  |  |  |  |  |  |  |  |  |
| --- | --- | --- | --- | --- | --- | --- | --- | --- | --- | --- | --- | --- |
|  |  |  |  | Mean distance |  |  |  | FWHM |  |  | TnC D159-R83 ionic binding |  |
| troponin | ligand |  | uP mean (std) | Δ | P |  | uP | Δ | P |  | uP | P |
| WT | apo |  | 30.18 (1.34) | -0.28 | 29.9 (0.9) |  | 2.99 | -0.61 | 2.38 |  | 0 | 0 |
| G159D | apo |  | 29.22 (0.98) | 0.35 | 29.59 (1.18) |  | 1.57 | -0.06 | 1.51 |  | 30062 | 28544 |
| G159D | EGCG |  | 31.17 (1.48) | 0.81 | 31.98 (1.59) | ? | 2.38 | -0.57 | 1.81 | ✓ | 8326 | 19890 |
| G159D | RVL |  | 32.13 (1.69) | -0.74 | 31.39 (1.51) | ✓ | 4.94 | -2.82 | 2.12 | ✓ | 18945 | 17269 |
| G159D | SilybinA |  | 30.09 (1.5) | 0.75 | 30.84 (1.04) |  | 4.4 | -1.97 | 2.43 | ? | 3101 | 15131 |
| G159D | SilybinB |  | 31.83 (1.83) | -1.11 | 30.72 (1.13) | ✓ | 1.9 | 0.74 | 2.64 | ? | 6554 | 8506 |
| G159D | ECG |  | 30.2 (1.2) | 0 | 30.2 (1.4) | ◇ | 1.8 | 0.7 | 2.5 |  | 30750 | 32250 |

When the recoupling molecules SB and RVL were added to the G159D simulation, the increase in mean angle upon phosphorylation was restored toward wild-type (SB, +3.58°, RVL, +9.28° compared with WT, +1.65°). In contrast, adding silybin A or ECG to G159D did not reverse the hinge angle change and resembled untreated G159D (SA, -6.4°, ECG -8.3° compared with apoG159D. -5.3°). The effect of EGCG on G159D hinge angle was anomalous: the mean angle decreased by 7.0° despite EGCG being a recoupling molecule. However, The EGCG angle distribution is bimodal or tri-modal suggesting multiple actions. If just the prominent peak centred on 103° is considered, EGCG may behave like the other recouplers. EGCG is both a recoupler and a desensitiser of actomyosin and this second property, not shared with SB or RVL, may be responsible for the occupancy below 80°. This is supported by the observation that the distribution plot for ECG which is a pure desensitizer is simpler and lacks the 80° component (Supplementary Information 2).

It is also notable that phosphorylation and mutations affect the broadness of the distribution of hinge angles which is reduced on phosphorylation of wild-type but not with G159D. G159D is substantially more rigid than wild type and it was proposed by Yang et al. that the strong bond between TnC R83 and D159 across the hinge played a part here (18). It is thus noteworthy that this interaction is substantially weakened by all the ligands binding even though there is no evidence for any ligand binding close to this site, implying an allosteric effect (Table 2A).

#### 2.3.4 Binding of small molecules to troponin

The molecular dynamics simulations allowed us to determine how the small molecules interact with G159D troponin and how this modifies the response to phosphorylation.

Figure 4 illustrates the preferred solution conformation of the ligands considered in this paper. silybin B and A are approximately flat; they are stereoisomers with the C ring in opposite orientations. The A, C, and D rings of EGCG have considerable homology with silybin but the orientation of the B ring gives the molecule a propellor-like shape. Resveratrol likewise has some structural homology with silybin and EGCG.

**Figure 4.**
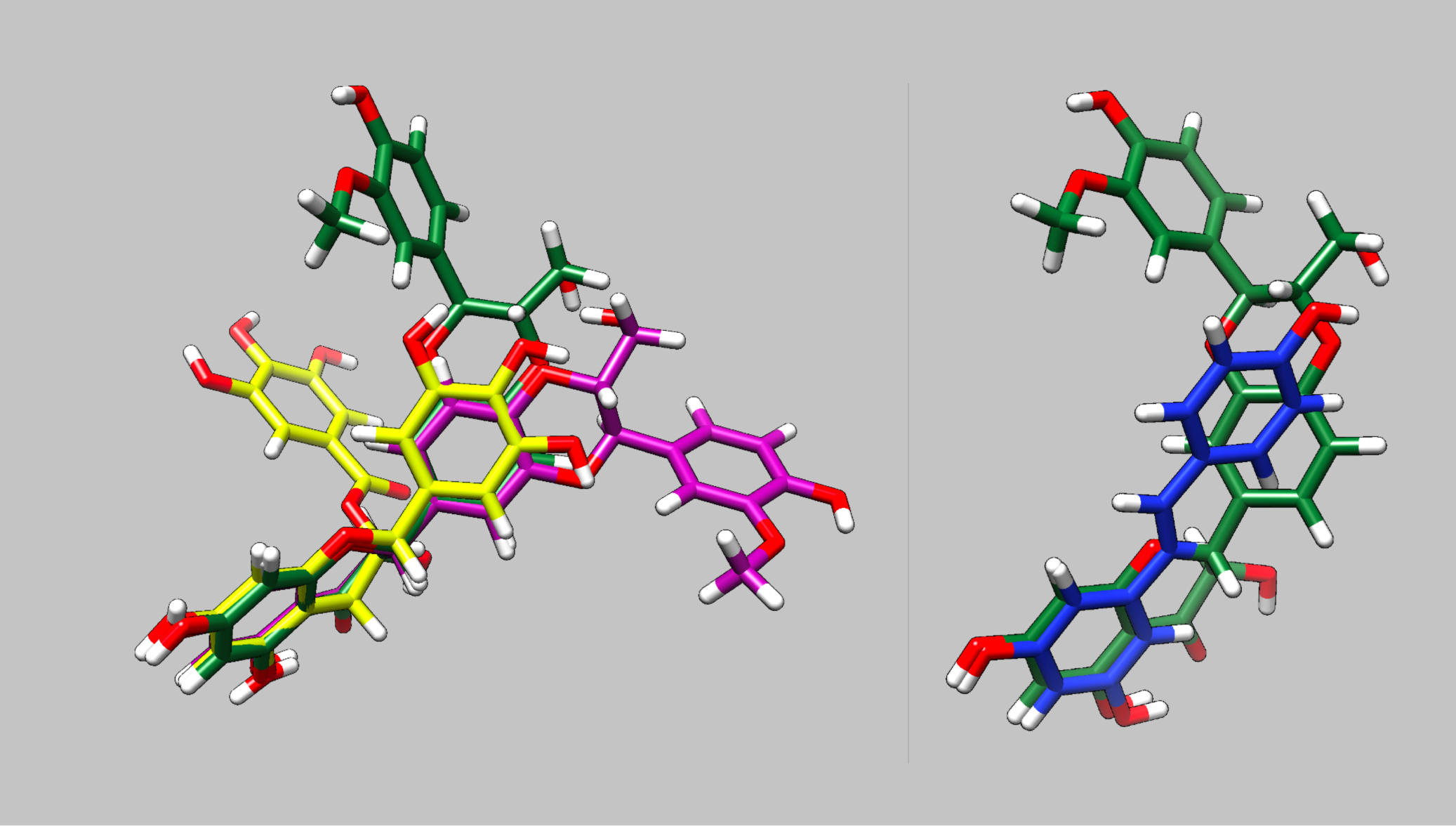
Parameterised solution configurations. silybin A (magenta) silybin B (green), EGCG (yellow) and resveratrol (blue). A and D rings of silybin A and B and A and C rings of EGCG are aligned. See Figure 1 and supplementary information 6 for identification of the atoms and rings.

##### 2.3.4.1 Binding energy

MMPBSA was used to calculate ΔG for ligand binding (Table 2B). ΔG was in the range 25.5-26.2 kJ/mole for EGCG, ECG, silybin A, and silybin B. This corresponds to a dissociation constant of around 6nM. ΔG for resveratrol appeared to be lower, consistent with its radically different binding footprint on troponin compared with the other recouplers. For binding to G159D, ΔG did not change much with phosphorylation except for silybin A where binding became significantly weaker on phosphorylation.

**TABLE 2B.** MMPBSA metrics.

| MMPBSA metrics |  |  |  |  |  |  |  |  |  |  |  |  |
| --- | --- | --- | --- | --- | --- | --- | --- | --- | --- | --- | --- | --- |
|  |  |  |  | MMPBSA Switch |  |  |  | MMPBSA Ligand binding |  |  | Fraction of time attached |  |
| Troponin | ligand |  | uP mean (std) | Δ | P |  | uP | Δ | P |  | uP | P |
| WT | apo |  | -108.43 (24.05) | -3.46 | -111.89 (13.91) |  |  |  |  |  |  |  |
| G159D | apo |  | -113.47 (13.75) | -3.53 | -117.0 (15.01) |  |  |  |  |  |  |  |
| G159D | EGCG |  | -109.26 (13.73) | -13.6 | -122.87 (13.56) |  | -26.12 (11.19) | -0.71 | -25.41 (10.14) |  | 0.93 | 0.88 |
| G159D | RVL |  | -104.91 (15.69) | -2.02 | -106.93 (15.23) |  | -13.71 (8.46) | -1.36 | -12.35 (7.56) |  | 0.49 | 0.33 |
| G159D | SilybinA |  | -113.65 (15.03) | 1.8 | -111.85 (16.08) |  | -25.48 (9.21) | -6.36 | -19.12 (9.26) |  | 0.95 | 0.82 |
| G159D | SilybinB |  | -106.72 (17.92) | 0.07 | -106.79 (13.52) |  | -26.16 (10.04) | 0.47 | -26.63 (10.69) |  | 0.95 | 0.88 |
| G159D | ECG |  | -108.1 (15.2) | -4.9 | -113.0 (16.6) |  | -25 | 0.9 | -25.9 |  | 0.92 |  |

##### 2.3.4.2 Ligand binding hotspots on troponin

We identified the most probable contacts on troponin using the CCPTraj procedures. The plot in Supplementary Information 8A shows the probability of interactions between the ligand and each amino acid of the troponin core. Although it is quantitative, the plot is one-dimensional and gives no idea of the stereochemistry of the interactions. In every case, discrete contact patterns were observed in unphosphorylated G159D that were different for each of the four small molecules and that changed upon phosphorylation as would be expected if they influence the phosphorylation-dependent structural and dynamic changes we have demonstrated.

One way of visualizing the interaction hotspots is to plot all contacts above a threshold (in this case interactions present >10% of the time) on representative models of the troponin core structure, see Supplementary Information 8B and movies, Supplementary Information 8C. Taken together these plots show that ligand binding is phosphorylation-dependent. In general, hotspots tended to be more dispersed after phosphorylation indicating that ligands explore a wider range of interactions compared with unphosphorylated although overall ΔG was not changed. Many of the phosphorylation-dependent hotspots are near the NcTnI-NctnC interface or the interdomain interface. It is also apparent that ligand interaction can change the conformation of the troponin core, consistent with the angle changes discussed above. Apparent contacts with the tail of troponin (around the TnI helix 1-helix 2 turns and the linker before TnT helix1) are likely to be artefactual since these parts of troponin will be in contact with the rest of the thin filament *in vivo* (22) (23).

In these plots there appear to be multiple discrete hotspots; an example of this is silybin B binding to unphosphorylated G159D where the hotspot appears to extend over a large area of the NcTnC surface which would not be possible for a single site given the size of silybin B and, in EGCG binding to unphosphorylated G159D, there are two hotspots on opposite sides of the troponin again not possible to be occupied by EGCG simultaneously. This is most likely due to the time-averaged nature of the CPTraj calculations whereas, in reality, ligand binding is a dynamic process. The ligand may occupy a number of locations at different times all of which appear in the model.

In order to understand ligand binding it is necessary to study it in 4 dimensions-this can only be done by directly observing the binding trajectories (Figure 5 and Supplementary Information 9). Observation confirms the stochastic nature of ligand binding. A ligand can bind at several discrete locations as was suggested for silybin B and EGCG. In the five 1500 ns runs, the initial locations can be different. Often the ligand goes to one location and stays there for prolonged periods, however, the ligand can move during a run, either by ‘creeping’ across the troponin or more often by dissociation and rebinding. In some cases, the ligand is free for a substantial part of the simulation, most notably with resveratrol (See Table 2B).

**Figure 5.**
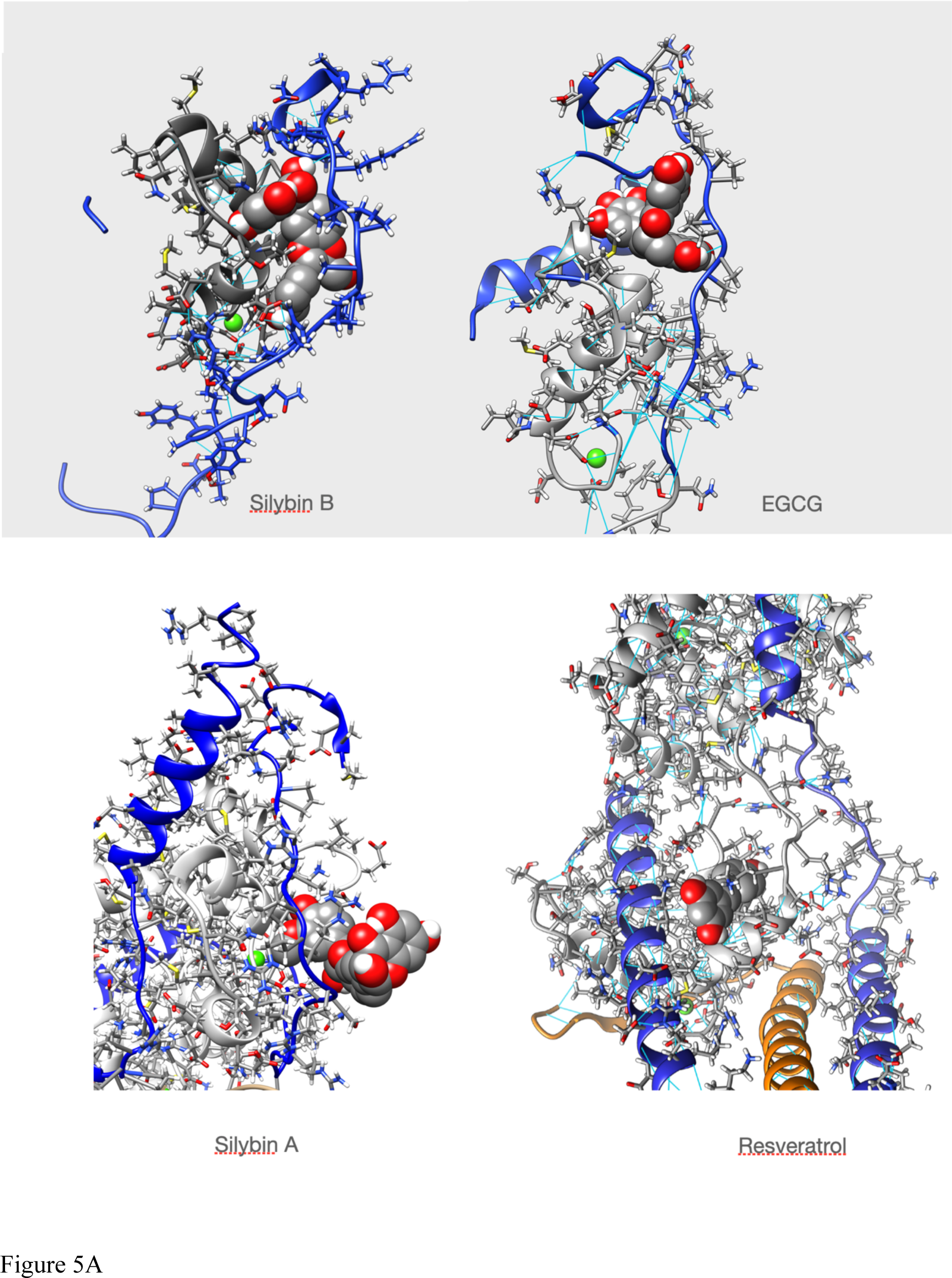
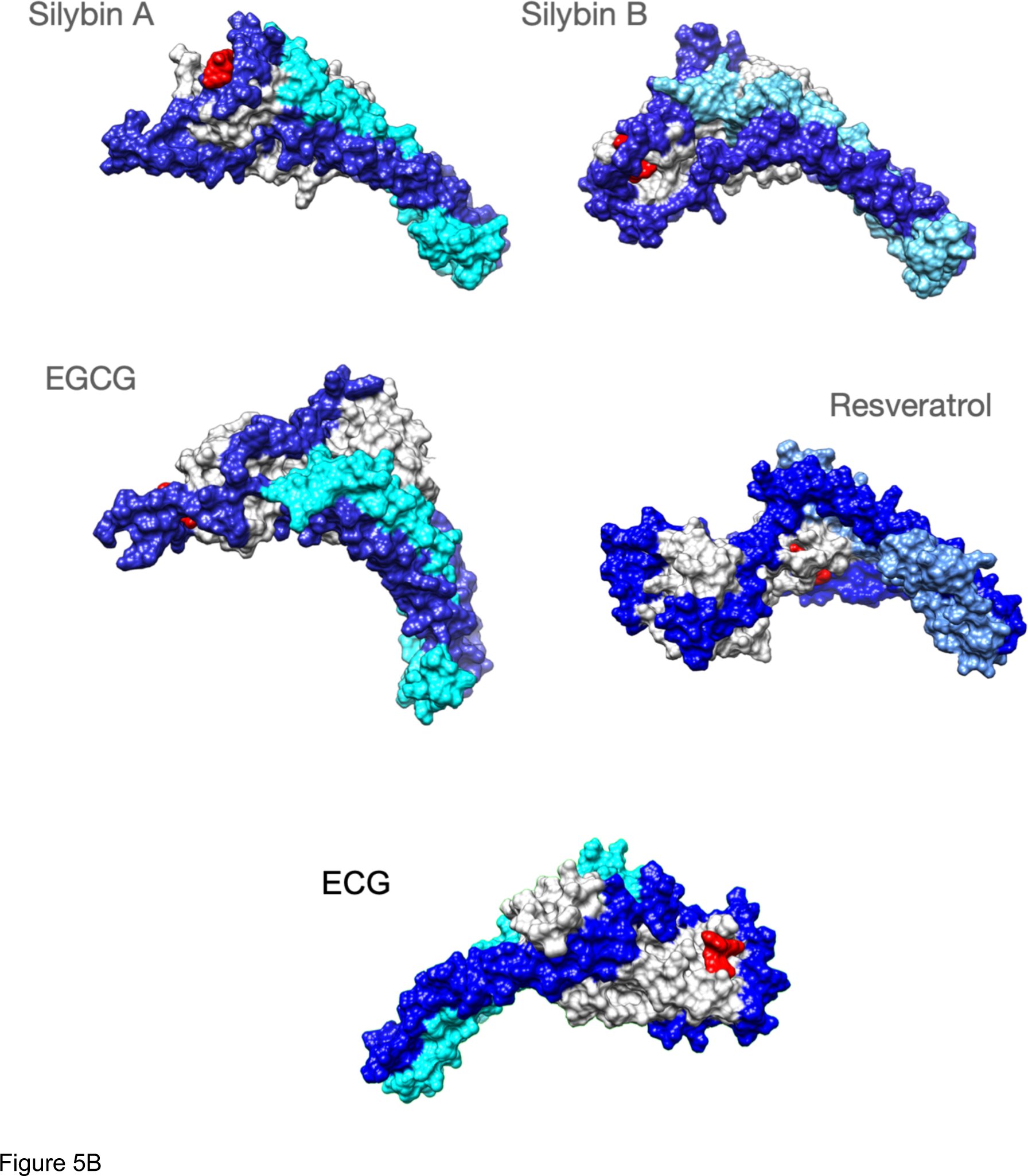
Ligands bound to unphosphorylated G!59D troponin in their most probable state. A-stick representation of the ligand interacting with TnC( light grey) and TnI(blue). TnT is light brown. Ligand is red The molecule is orientated to give the best view of the interaction. B-Whole troponin surface representations. TnI is blue, TnC is grey and TnT is light blue or cyan. Ligand is red. Single frames are selected from the full trajectory (3750 frames) to illustrate the most common positions of the ligands. Models are orientated to show the ligand binding optimally. Full trajectories are illustrated in supplementary information 9.

We searched for a pattern of contacts that was associated with recoupling. Supplementary Information 9 and Figure 5 show the analysis of selected whole trajectories and highlights those positions where the ligand is often found. Observation suggests for silybin B there are two common positions: at the end of cTroponin, intercalated between the NcTnI peptide and NcTnC A and B helices and at the ‘top’ of the hinge, interacting with CcTnT and the ‘inhibitory peptide’ (137-148). SilybinB is bound nearly all the time to unphosphorylated G159D and tends to go to one site or the other. Therefore the time average in Supplementary information 8B is overwhelmingly a mixture of two separate hotspots. Figure 5 shows a frame from one run where SilybinB occupies site A for most of the 1500ns run. When phosphorylated (SEP), silybin B is almost never at site A. It is often in parts of the tail and CcTnC and sometimes also at top of hinge like SilybinB site B.

Silybin A also binds primarily near the N terminus of TnI and the helix A-B loop of EF I (Figure 5 and Supplementary Information 9). Unlike Silybin B Silybin A does not appear to be intercalated between NcTnI and NcTnC but sits on the surface with only the A ring engaged. A secondary site is at the ‘top’ of the interdomain hinge region. All of these contacts are lost on phosphorylation.

In unphosphorylated G159D, EGCG occupies the site intercalated between NcTnI and NcTnC (site A) identified in silybin B part of the time, although it does not use the same set of contacts and there are additional hydrogen bonding to the switch peptide, perhaps associated with its desensitising property (Figure 5 and Supplementary Information 9). None of these sites are occupied when troponin is phosphorylated.

In contrast ECG primarily occupies a site on the opposite side of NcTnC where it sits on the surface approximately flat with the B ring extending away from troponin (Supplementary Information 2).

Resveratrol binds less frequently to troponin than the others ( it was free for 45% of the MD runs see Table 2B) and most often makes contacts with a completely different region of troponin, inserted into the C-terminal domain of troponin C helix E, close to the interdomain interface (Figure 5, Supplementary Information 8 and 9).

## 3 DISCUSSION

### 3.1 Silybin B, EGCG and resveratrol can restore lusitropy to mutant myocytes that have impaired lusitropy

Lusitropy, the increase in the rate of cardiac muscle relaxation, is an essential component of the heart’s response to adrenergic activation (5). It is predominantly mediated by PKA phosphorylation of cardiac troponin I (24,25). However in genetic cardiomyopathies due to mutations in thin filament proteins, lusitropy is much reduced and *in vitro* PKA phosphorylation does not alter Ca^2+^-sensitivity, the usual measure of the effect of phosphorylation (5,11). Basic and animal studies indicate that specific suppression of lusitropy can induce symptoms of heart failure under stress and clinical studies have shown that lack of response to adrenergic activation is associated with increased adverse cardiac events (13) (5). Thus suppression of lusitropy by mutations may be a significant disease mechanism for cardiomyopathy.

Restoration of the lusitropic response is therefore a suitable target for therapy. We have investigated a series of small molecules, acting on troponin, that can restore the phosphorylation-dependent modulation of Ca^2+^-sensitivity (recoupling) at the single myofilament level as potential treatments (16) (15). These molecules appeared to be promising since they all gave total restoration *in vitro* that was independent of the mutation that caused the dysfunction in lusitropy (Summarised in Supplementary Information 1). However, it was not established whether these small molecules had any effect on intact muscle and the molecular mechanism by which many different small molecules could cause a single outcome, recoupling, was unknown.

In this study, we have addressed these questions for five small molecules with a range of functions and structures. Silybin B and resveratrol are structurally distinct yet both have a pure recoupling activity *in vitro*, whilst EGCG is bifunctional, being both a recoupler and a desensitiser. As negative controls, we studied silybin A, the stereoisomer of silybin B, and ECG, a close analogue of EGCG, that have no recoupling activity i*n vitro*.

We tested these small molecules for their ability to restore lusitropy in two previously studied mutant systems. Myocytes from ACTC E99K mice and guinea-pig myocytes transfected with the TNNT2 R92Q mutation were challenged with the β1 specific adrenergic agent dobutamine (20,21). The results correlated with *in vitro* results: the mutant myocytes had suppressed lusitropy but wild-type lusitropy could be restored by silybin B, EGCG and Resveratrol but not by silybin A. This result confirms that recouplers studied *in vitro* do restore lusitropy, meaning that these small molecules have therapeutic potential.

### 3.2 Silybin B, EGCG, and resveratrol can restore the dynamics of the G159D mutant troponin core to wild-type values

To understand the molecular mechanism by which silybin B, EGCG, and resveratrol restore lusitropy we extended our molecular dynamics studies of the structure and dynamics of the troponin core that have defined how phosphorylation at cTnI Ser22 and 23 modulates the dynamics of troponin and how the DCM-associated mutation, G159D interferes with this modulation (18,26). Several phosphorylation-dependent metrics may be derived from the MD simulations; the most useful for this study are the change in the distribution of the angle between the A and B helices of cTnC ( a measure of the opening and closing of the hydrophobic patch) and changes in the hinge angle between the two quasi rigid domains of troponin ( NcTnC and cTnI 1-33 vs CcTnC / TnT / TnI helices H1 and H2 see (27)).

Phosphorylation, in wild-type, causes an increase in the mean A/B helix angle and a reduction in the range of motion (FWHM); phosphorylation decreases the mean hinge angle whilst also reducing the FWHM parameter.

In contrast, phosphorylation of G159D troponin has an opposite effect on most of the metrics compared with wild-type. The G159D mutant troponin has an intrinsically lower A/B helix angle and FWHM than wild-type and phosphorylation causes an increased mean helix A/B angle whilst there is a decrease in interdomain hinge angle when G159D is phosphorylated. We have proposed that these abnormalities seen in G159D dynamics are causative of the uncoupling effect of the mutation and therefore could be used to test for recoupling.

In the presence of the three recoupling compounds the parameters for A/B helix angle and the interdomain hinge angle of G159D troponin are largely restored to wild-type values (See Figure 6 for a summary), thus recoupling involves restoration of wild-type dynamic behaviour in troponin at the atomic level. It is perhaps relevant that the strong ionic and H bonding between the mutant amino acid cTnC D159 and T83 across the interdomain interface thought to be responsible for the reduced range of hinge motion in the mutant, is much reduced by ligand binding. It has been noted that ligands may occupy multiple sites only one of which may be relevant to recoupling and also that ligand is not always bound, especially with resveratrol (Table 2B). These factors may obscure the full effects of the ligands on the MD metrics.

**Figure 6.**
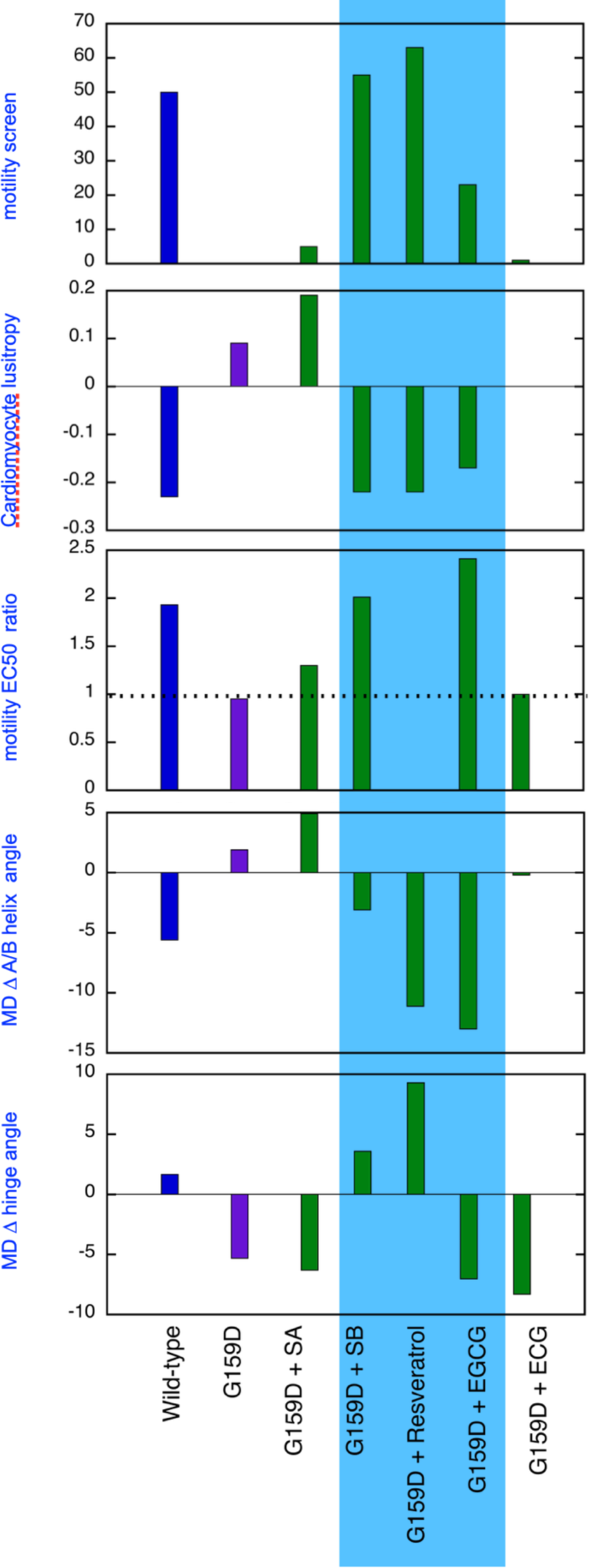
Comparison of the effect of phosphorylation of cardiac troponin on its biochemical, physiological and molecular dynamics parameters, its suppression by the TnC G159D DCM related mutation and its restoration by small molecules. Parameters compared are lusitropy in intact myocytes (from Fig 3), Fixed [Ca2+] screen for coupling in thin filaments measured by IVMA from Sheehan et al., [22] and effect of phosphorylation on EC_50_, (EC_50_P/EC_50_uP) in thin filaments measured by IVMA (from Figure 1), Change on A/B angle on phosphorylation and change in hinge angle upon phosphorylation from Molecular dynamics simulations (from Figure 2, Tables 1A and 1B). The recouplers are highlighted in blue.

Importantly, silybin A and ECG that do not recouple do not alter the phosphorylation-dependent metrics of G159D significantly, confirming the specific effect of the recouplers. It is notable that silybin B, EGCG, and resveratrol affect the parameters for unphosphorylated G159D with little effect on phosphorylated G159D in parallel with the i*n vitro* motility measurements that also show that G159D behaves like phosphorylated wild-type independent of phosphorylation and the small molecules primarily alter the properties of unphosphorylated troponin (Figure 1 and Supplementary Information 4).

### 3.3 Ligand binding and recoupling

Understanding how the recouplers function cannot be precise because of the disordered nature of troponin in the regions where the ligands contact and the weak, stochastic binding of the ligands. Nevertheless, the ligands do satisfy the basic criterion of allosteric effectors since they bind differently to unphosphorylated and phosphorylated G159D troponin. There is no significant change in binding energetics of the ligands to unphosphorylated and phosphorylated troponin, instead we observe that ligands are concentrated on a few ‘hotspots’ in unphosphorylated troponin, that are not occupied in phosphorylated troponin and that the ‘hotspots’ are more dispersed in phosphorylated troponin. We have concentrated our analysis on the effects of the ligands on unphosphorylated G159D troponin. The most common poses of bound silybin B and EGCG are found to be intercalated between the disordered N terminus of cTnI, adjacent to the phosphorylatable serines 22 and 23, and the helix A and B region of TnC. Since these poses are only seen in unphosphorylated troponin it is feasible that their occupancy is the cause of recoupling. Compatible with this hypothesis is the observation that silybin A binds to roughly the same region of NcTnI but is not intercalated between TnI and TnC, presumably because the stereoisomer does not fit the space. Similarly, ECG binds predominanatly to the surface of cTnC at a site only rarely occupied by EGCG.

Our study included resveratrol since it was functionally identical to silybin B in single filament and myocyte assays but had a radically different chemical structure. In 45% of the MD runs resveratrol is unbound (Table 2B) and its contacts with unphosphorylated troponin are also very different from silybin B and EGCG. In unphosphorylated troponin it appears that resveratrol is mostly located near helix E in the C terminal domain of cTnC near the hinge interface, possibly intercalated between the two domains. As with the other recouplers, binding is reduced and dispersed on phosphorylation. Therefore, we cannot propose a single mechanism for recoupling; a study of a wider range of recouplers would be needed to attempt to resolve the question.

### 3.4 CONCLUSIONS

A primary aim of this study was to demonstrate that the uncoupling phenomenon caused by cardiomyopathy-related mutations and the ability of small molecules to restore coupling, first found with pure thin filaments *in vitro*, is also relevant at the cellular and atomistic levels. Figure 6 summarises our results that support this proposition. Uncoupling is manifested at the myocyte level as an insensitivity of relaxation rate to β adrenergic stimulation and at the atomistic level as the failure of phosphorylation to increase interdomain angle or decrease A/B angle. It is evident that silybin B, resveratrol, and EGCG restore the native parameters to the mutant troponin whilst silybin A and ECG do not, as predicted.

We have shown Silybin B has the potential to play a role in safeguarding the heart against cardiomyopathy and Silybin A does not compromise this activity (16).

Racemic Silybin is a readily available and safe nutraceutical, as is resveratrol (28). Further research on these compounds as treatments to restore lusitropy is thus indicated.

The compounds studied here are all polyphenols that may have multiple pharmacological activities that render them unsuitable for specific treatments (29-31). However, at least 25 small molecules have been shown to be recouplers. Further work on these *in silico* could define the specific properties needed for recoupling and allow for the prediction and design of new molecules that recouple without additional activities that could be used in the treatment of genetic cardiomyopathies.

## METHODS

4.1 Reagents were obtained from Sigma-Aldrich, except for silybin A and silybin B that were prepared by Prof Vladimir Kren and Dr David Biedermann (32).

4.2 The movement of synthetic thin filaments containing G159D troponin C over immobilised myosin was measured by *in vitro* motility assay as described by (19). The fraction motile parameter is plotted here. We measured Ca^2+^ activation curves for wild type and G159D troponin-containing thin filaments in the unphosphorylated and phosphorylated states obtained by phosphatase and kinase treatments. A single Ca^2+^-concentration assay was used as a rapid screen for coupling and recoupling activity. [Ca2+] was 0.1µM and [ligand] was 100uM; the method is described in Supplementary Information 3.

4.3 Mouse cardiomyocyte isolation protocol and measurement of cell shortening was based on previously described methods using the Ionoptix system (Ionoptix, Milton, MA). Cell contraction was measured using a charge-coupled device video camera (MyoCam-S, IonOptix) connected to a personal computer running IonWizard software (33). Guinea pig myocyte contractility was measured in the Cytocypher instrument as described by (34) using transfected guinea-pig myocytes prepared as described by (21). Incubation medium was Krebs-Hensleit buffer with added 1mM CaCl_2._ Contractility in each dish of myocytes was measured in the absence and then presence of 0.4µM dobutamine and 50nM ICI 118,551.

### 4.4 Molecular Dynamics Studies

*4.4.1 MD simulations of troponin* were performed as described by Yang et al (18). Parameterisation of the ligands in this study was based on a general AMBER force field (GAFF) which is suitable for small organic molecules (35). Ligand partial charges were derived from the ground state structures which were found by following conformational search, geometry optimisation, and thermal correction procedures, as detailed below: An ensemble of conformers were generated for each ligand using the ETKDG algorithm as implemented by RDKit. (36)

Each conformer was optimised to a local minimum. Each optimised structure was subjected to thermal correction with frequency to calculate the Gibbs free energy. (37-39). The structure with the lowest ground-state energy was picked as the ground-state structure

From the ground state structures, restrained electrostatic potential (RESP) calculations were used to obtain the charge distribution on the molecule (40). Gaussian16 (41) was used for both geometry optimisation and RESP calculation. Geometry optimisation was done with B3LYP functional with cc-pVDZ basis set and the polarisable continuum model (PCM) in water as an implicit solvent (42-45). RESP calculation was done with Hartree-Fock method and 6-31G* basis set in vacuum. (46). AmberTools was then used to generate the charge parameters for each atom.

*4.4.2 Analysis* Representative structures, binding energy estimation by MMBPSA, interhelical angle distribution and hinge angle analysis were performed as described by (18). CPPTraj was used to determine the contact/interactions. Pytraj (47), a Python package binding to cppraj program (48), was used for the distance measurements in this work. The ligand was deemed to be in contact with a residue when the minimal distance between any atom of the ligand and a residue was lower than 2.5 Å. The same was applied to ligand atom interactions where a ligand atom was deemed to be in contact with the protein when the minimal distance between atoms of the ligand and any part of the protein was lower than 2.5 Å.

## ACKNOWLEDGEMENT

SM was supported by British Heart Foundation programme grant (RG/ 11/20/29266).

AS has a Ph.D. studentship funded by the BHF Centre of Research Excellence (RE/13/4/30184).

ZY is supported by an awarded Ph.D. studentship from the Wellcome Trust grant number 108908/Z/15/Z A 4-year PhD in theoretical systems biology and bioinformatics.

VK is supported by the project of the Ministry of Education, Sports, and Youth of the Czech Republic (MEYS) “Talking microbes-understanding microbial interactions within One Health framework” (CZ.02.01.01/00/22_008/0004597)

CR and AS are supported by the British Heart Foundation Grant PG/18/68/33883.

## Notes

### Competing Interest Statement

The authors have declared no competing interest.

### Summary of Updates

Revised for clarity and typos following referee reports. Reformatted for JBC

