## Supplementary material for "Nutraceuticals Silybin B, resveratrol and epigallocatechin-3 gallate (EGCG) bind to cardiac muscle troponin to restore the loss of lusitropy caused by cardiomyopathy mutations *in vitro, in vivo*, and *in silico*"

**CONTENTS:**

**1** Many compounds re-couple many mutations.

**2** Comparison of ECG and EGCG in vitro, ex vivo and in silico.

**3** Single [Ca2+] screen for recoupling

**4** Effects of TnI PKA phosphorylation on $EC_{50}$ of wild-type and TPM1 G159D mutant thin filaments and its modulation by small molecules.

**5** Effects of dobutamine on cardiac myocyte contractility.

**6** Lusitropy and the effect of small molecules measured in cardiomyocytes

**7** Preferred structure of small molecules with atoms and rings labelled.

**8** CCPtraj analysis of ligand bindin, ligand hotspots on representative structures, link to movies

**9** snapshots from single 1500ns MD trajectories, 7500 total frames

**SUPPLEMENT 1**

Many compounds re-couple many mutations.
Full set of those tested.

| | Egcg | SA | SB | resveratrol | DHSA | DHSB | Silybin | Blagg 7 | Blagg 11 |
|---|---|---|---|---|---|---|---|---|---|
| **TPM1 R180G** | + | + | + | + | + | + | + | + | + |
| **TNNC1 G159D** | + | + | + | | + | + | | | |
| **ACTC E99K** | + | ++ | ++ | ++ | | | + | + | |
| **TPM1 E54K** | + | | | | | | + | + | + |
| **MYBPC 3 R820W** | + | | | | | | | | |
| **TNNT2 R92Q** | ++ | ++ | + | + | | | | | |
| **TNNT2 K280N** | + | | | | | | | | |
| **TNNT2 Δ28** | + | | | | | | | | |
| **TNNT2 K2732N** | + | | | | | | | | |
| **TNNT2 S179F** | + | | | | | | | | |
| **TNNT2 ΔE160** | + | | | | | | | | |
| **ACTC E361G** | +* | | +* | | | | | | |
| | | Black: IVMA<br>Red: IVMA and myocyte<br>Green: not recoupler<br>Purple: reverse recoupler<br>*Tested in papillary muscle | | | | | | | |

Blank means not tested; we found no compound that worked with one mutant and not another.

**SUPPLEMENT 2 Comparison of ECG and EGCG in vitro, ex vivo and in silico.**

A ECG differs from ECGC by removal of one phenoloic OH from the B ring

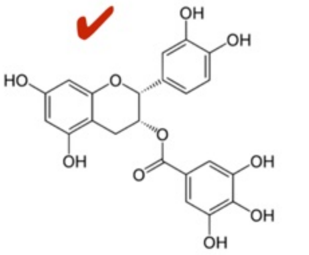

Epicatechin Gallate (ECG)

B Both ECG and EGCG are desensitisers in skinned muscle fibres (Tadano, N. *et al.* Biological actions of green tea catechins on cardiac troponin C. *British journal of pharmacology* **161**, 1034–1043 (2010).)

C ECG is a pure desensitiser in the in vitro motility assay (Sheehan, A. *et al.* Molecular Defects in Cardiac Myofilament Ca2+-Regulation Due to Cardiomyopathy-Linked Mutations Can Be Reversed by Small Molecules Binding to Troponin. *Frontiers in physiology* **9**, 300–12 (2018).).

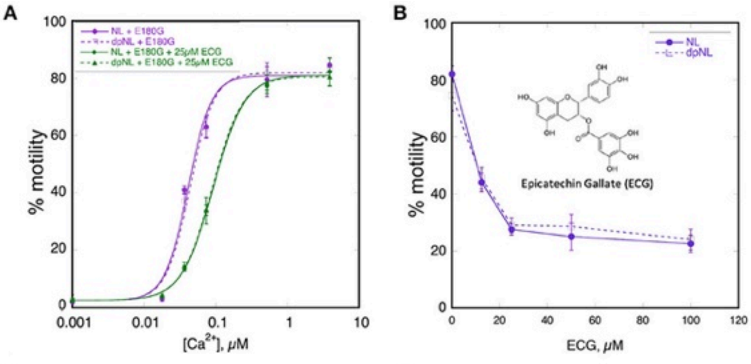

D ECG, unlike EGCG, does not restore A/B angle or hinge angle It also does not restore interdomain distance change or reduce the D159>R83 interaction.

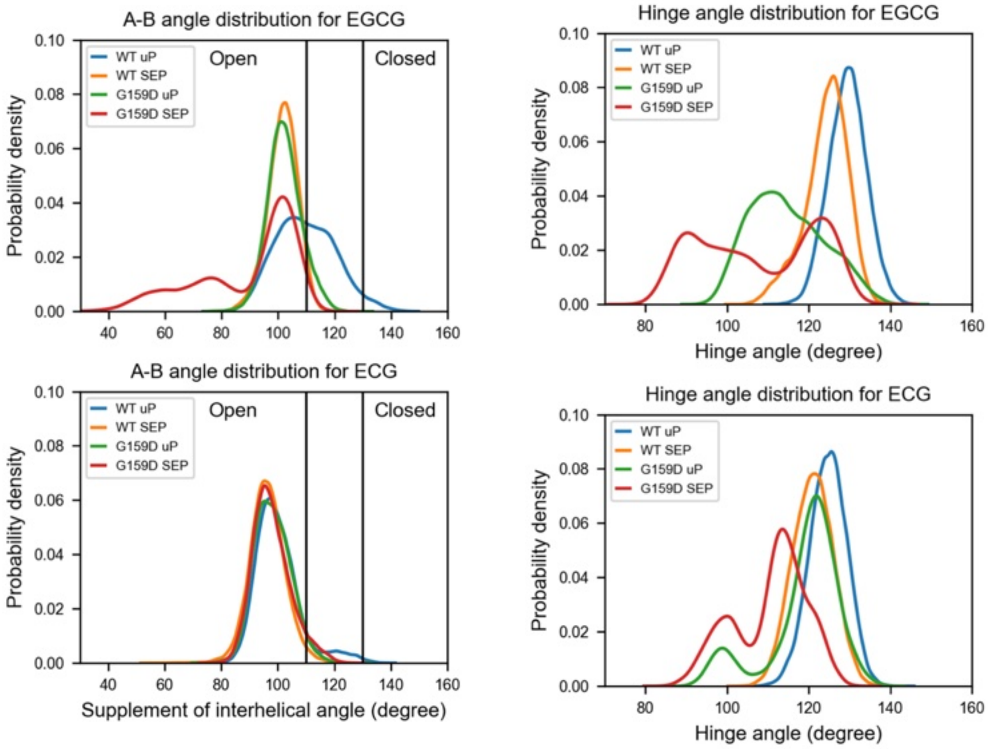

A/B angle: ECG loses the bimodal features of EGCG (both wt and G159D), compatible with its single (desensitising) function
Hinge angle : EGCG loses peaks at 90 in SEP and uP, peaks around 120 dominate and peaks around100-110 diminished ,
especially in uP. In effect a reduction in complexity, compatible with loss of the recoupling function. It would be interesting to see if the hotspots are selectively diminished also.

E ECG bound to troponin: snapshots from 75000 frames, run 02

**ECG Run 02 623** **ECG Run 02 1788** **ECG Run 02 2638** **ECG Run 02 3579**

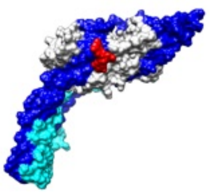
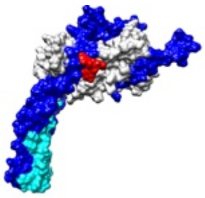
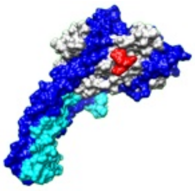
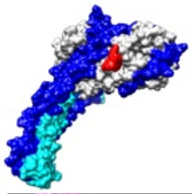

H12,H13 _____glu 10 OE1

EGCG

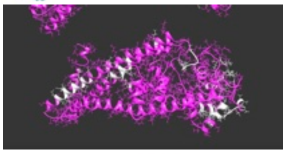

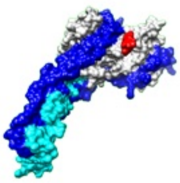
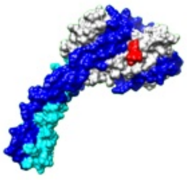
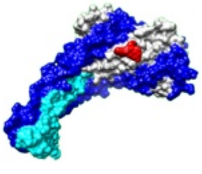
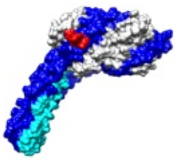

**ECG Run 02 4469** **ECG Run 02 5436** **ECG Run 02 6335** **ECG Run 02 7280**

**SUPPLEMENT 3**

Explanation of the single point screen for ligand effects on coupling

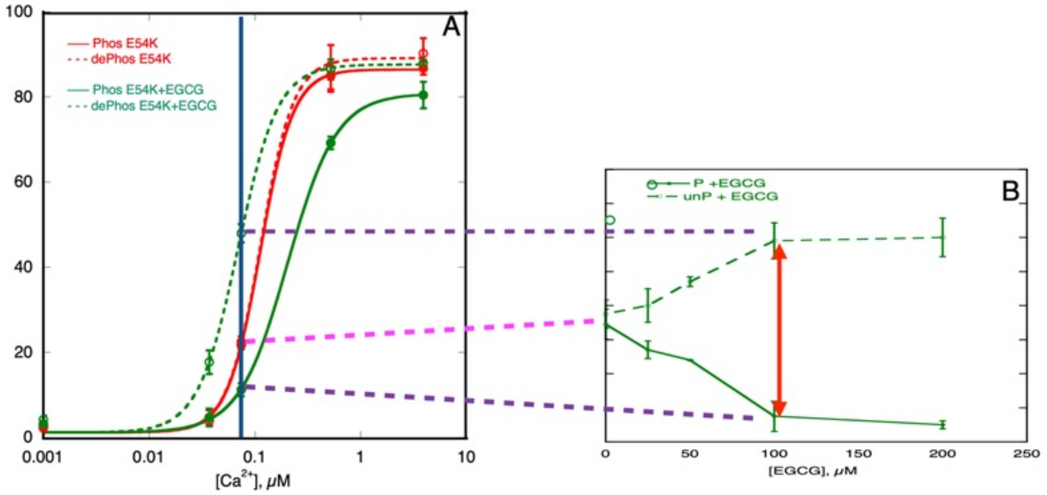

**Figure 2.4: Screening principle.** (A) The effects of EGCG on $Ca^{2+}$ regulation of E54K-containing thin filaments. (B) EGCG dose response data of thin filament motility measured at a constant $Ca^{2+}$ concentration corresponding to the $Ca^{2+}$ $EC_{50}$ for mutant filament mobility. Red lines with points represent mutation with no drug and Green lines with points represent mutation treatment with EGCG (100μM). Solid lines with points represent phosphorylated TnI and broken lines with points represent unphosphorylated TnI. Purple and pink broken lines illustrate the relationship between thin filament motility at a constant intermediate $Ca^{2+}$ concentration.

**SUPPLEMENT 4**

Effects of TnI PKA phosphorylation on $EC_{50}$ of wild-type and mutant ( TPM1 E180G and TNNC1 G159D) thin filaments and its modulation by small molecules. Reconstituted thin filaments measured by *in vitro* motility assay.

Pink,wild-type; blue, TPM1 E180G mutant; green, desensitised; ✓, phosphorylation and $EC_{50}$ are coupled or recoupled

| Mutation | EC50 of phosphorylated thin filament,<br>µM ± SEM | EC50 of unphosphorylated thin filament,<br>µM ± SEM | Ratio EC50 P/unP ± SEM |
|---|---|---|---|
| **Native Tm** | 0.0647 ± 0.0066 (2) | 0.0335 ± 0.0006 (2) | 1.93±0.23***✓ |
| **Native Tm +EGCG** | 0.26 ± 0.03 ** | 0.15 ± 0.02**†† | 1.73 ± 0.16 ✓ |
| **Native Tm + SA** | 0.0658 (1) | 0.0354 (1) | 1.86✓ |
| **Native Tm + SB** | 0.0545 (1) | 0.0302 (1) | 1.80✓ |
| **Native Tm + DHS A** | 0.0632 (1) | 0.0330 (1) | 1.91✓ |
| **Native Tm + DHS B** | 0.0660 (1) | 0.0331 (1) | 1.99✓ |
| **Tm E180G** | 0.0341± 0.0032 (9) | 0.0363 ± 0.0047 (9) | 0.95 ±0.025** |
| **Tm E180G +EGCG** | 0.11 ± 0.01* | 0.048 ± 0.003 *† | 2.41 ± 0.32 †✓ |
| **Tm E180G + SA** | 0.0654 (1) | 0.0490(1) | **1.33** |
| **Tm E180G + SB** | 0.0616 (1) | 0.0306 (1) | 2.01✓ |
| **Tm E180G + DHS A** | 0.0209 (1) | 0.0341 (1) | **0.61** |
| **Tm E180G + DHS B** | 0.0627 (1) | 0.0259 (1) | 2.42✓ |
| **TnC G159D** | 0.092±.004 (5) | 0.095±0.0005 (5) | 0.97 |
| **TnC G159D + EGCG** | 0.19±0.03 (5) | 0.088±0.005 (5) | 2.2✓ |
| **TnC G159D + Silybin A** | 0.141 | 0.136 | 1.0 |
| **TnC G159D + Silybin B** | 0.121 | 0.053 | 2.3✓ |

#### SUPPLEMENT 5

Effects of dobutamine on cardiac myocyte contractility.

| | n | Amplitude Raw Data, $\mu m$ | sem | $T_{90}$ contraction, sec | sem | $T_{90}$ relaxation, sec | sem |
|---|---|---|---|---|---|---|---|
| **mouse WT** | 57 | 3.566 | *2.427* | 0.046 | *0.014* | 0.168 | *0.052* |
| **mouse WT+ Dob** | 38 | 5.27 | *3.08* | 0.044 | *0.010* | 0.134 | *0.037* |
| **Mouse E99K** | 58 | 3.593 | *2.06* | 0.065 | *0.018* | 0.246 | *0.08* |
| **Mouse E99K+ Dob** | 43 | 5.195 | *3.0* | 0.066 | *0.02* | 0.277 | *0.086* |
| **GP WT** | 64 | 4.25 | *1.24* | 0.102 | *0.009* | 0.464 | *0.057* |
| **GP WT + Dob** | 45 | 4.12 | *1.43* | 0.0883 | *0.006* | 0.365 | *0.042* |
| **GP R92Q** | 45 | 3.0 | *0.86* | 0.98 | *0.019* | 0.466 | *0.077* |
| **GP R92Q + Dob** | 40 | 3.56 | *0.81* | 0.102 | *0.014* | 0.428 | *0.061* |

| ±dob student | mouse | e99k | GP | R92Q |
|---|---|---|---|---|
| **Amp, %** | 0.0024 | 0.0045 | 0.27 | 0.35 |
| **ttp90** | 0.628 | 0.863 | 0.28 | 0.88 |
| **ttb90** | <.0001 | 0.148 | 0.028 | 0.22 |

**SUPPLEMENT 6**

**Lusitropy and the effect of small molecules measured in cardiomyocytes**

| | | lusitropy | sem | student t | | cells/ hearts |
|---|---|---|---|---|---|---|
| **mouse** | WT | -0.20200 | 0.037000 | 0.00024000 | | 38/17 |
| | Mut | 0.12600 | 0.029000 | 0.0015000 | | 30/16 |
| | mut SB | -0.25400 | 0.044000 | 0.00020000 | | 8/3 |
| | mut Resv | -0.33100 | 0.047000 | 0.0010700 | | 8/3 |
| | mut SA | -0.038000 | 0.055000 | 0.54000 | | 6/2 |
| | mut EGCG | -0.22200 | 0.020000 | 0.00017000 | | 8/6 |
| **guinea pig** | WT | -0.23700 | 0.038000 | 0.00042000 | | 120/8 |
| | Mut | 0.091000 | 0.056000 | 0.18400 | | 85/5 |
| | mut SB | -0.13000 | 0.065000 | 0.0059000 | | 80/5 |
| | mut Resv | -0.17000 | 0.028000 | 0.050000 | | 64/4 |
| | mut SA | 0.18660 | 0.080000 | 0.14800 | | 50/3 |
| | mut EGCG | -0.17000 | 0.060000 | 0.054000 | | 51/3 |

**SUPPLEMENT 7**

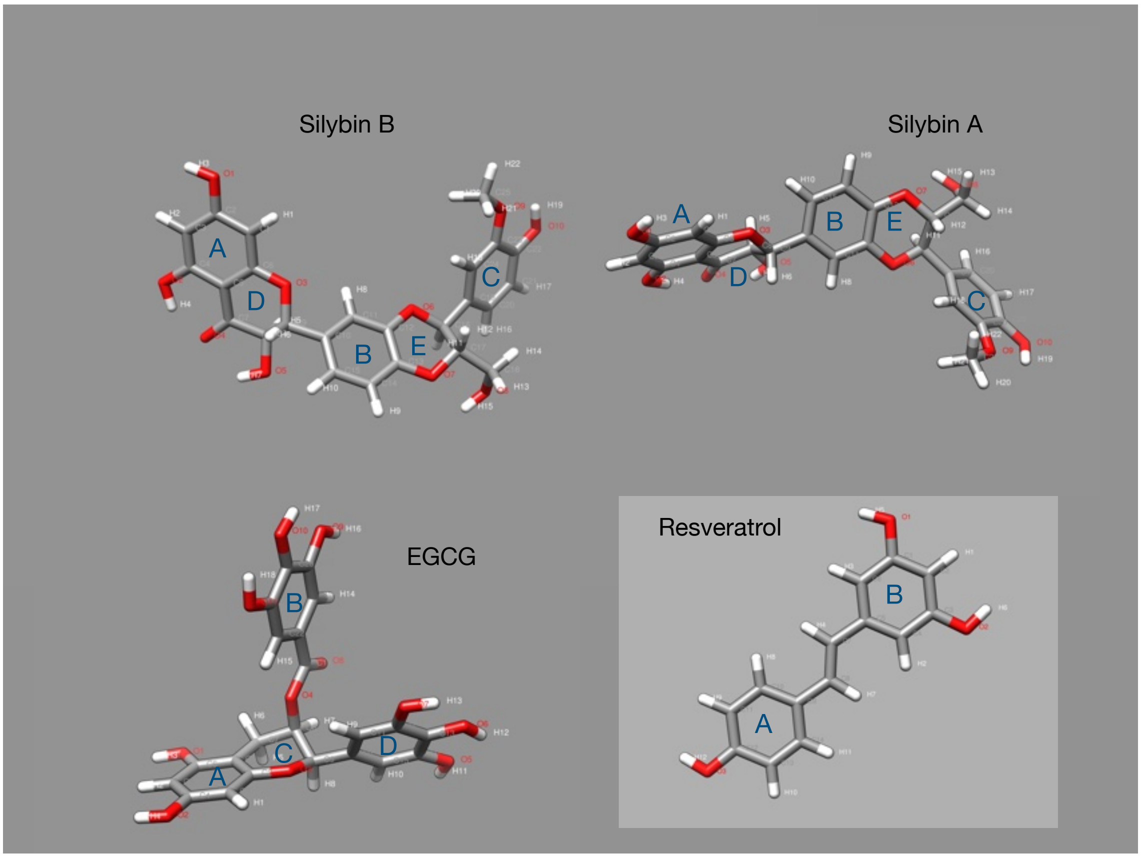

preferred structure of small molecules with atoms and rings labelled.

**SUPPLEMENT 8**

A CCPtraj analysis of ligand binding,

B ligand hotspots on representative structures,

C movies

8A

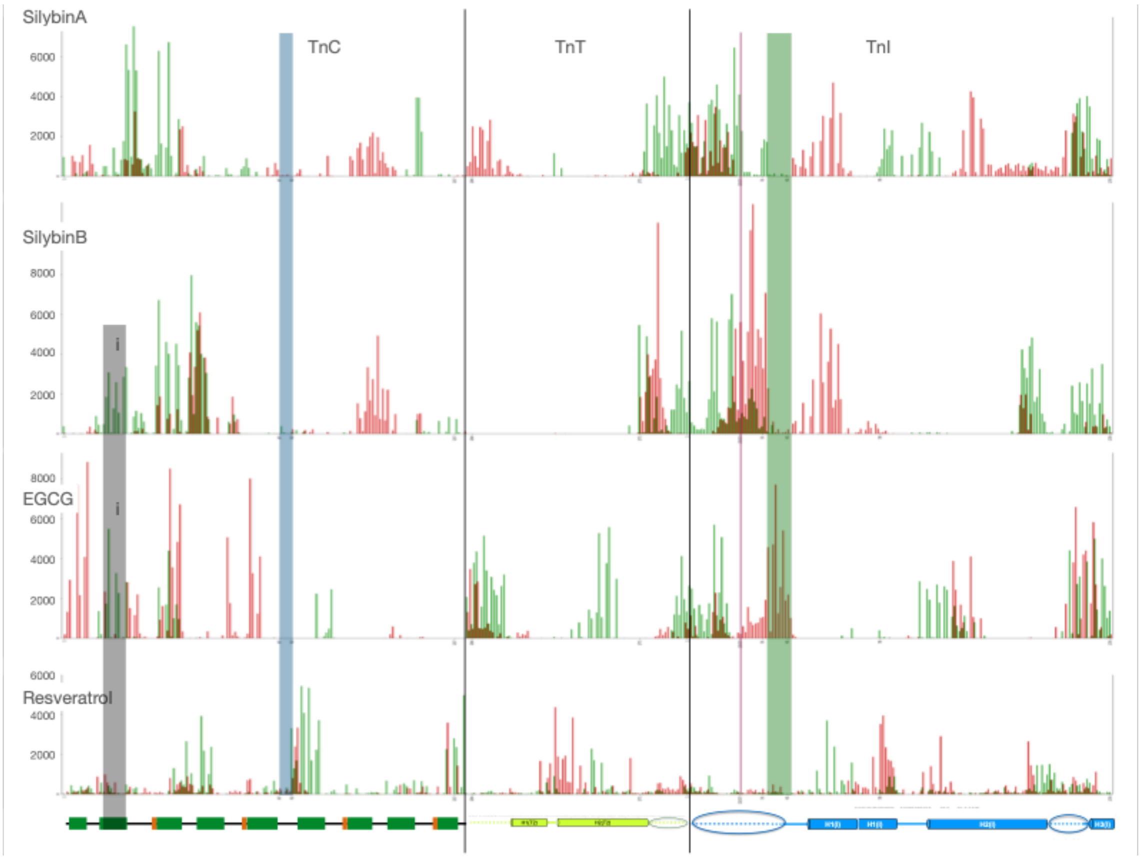

8B

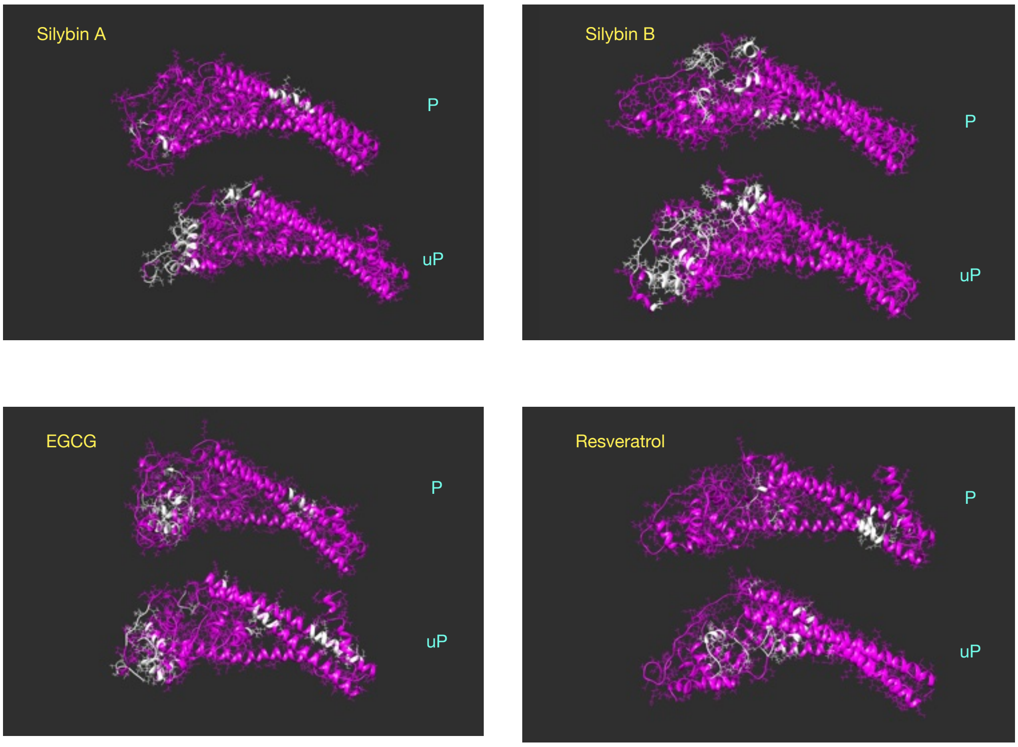

8C

For movies follow this link.

https://www.dropbox.com/scl/fo/zq4ttu6kpub3kcethaseu/AHBbV-mwDr6dsSJIr-mEV5A?rlkey=fshl5nivn68j8nrnjq6n2trgh&dl=0

**SUPPLEMENT 9**

snapshots from single 1500ns MD trajectories, 7500 total frames

Silybin B

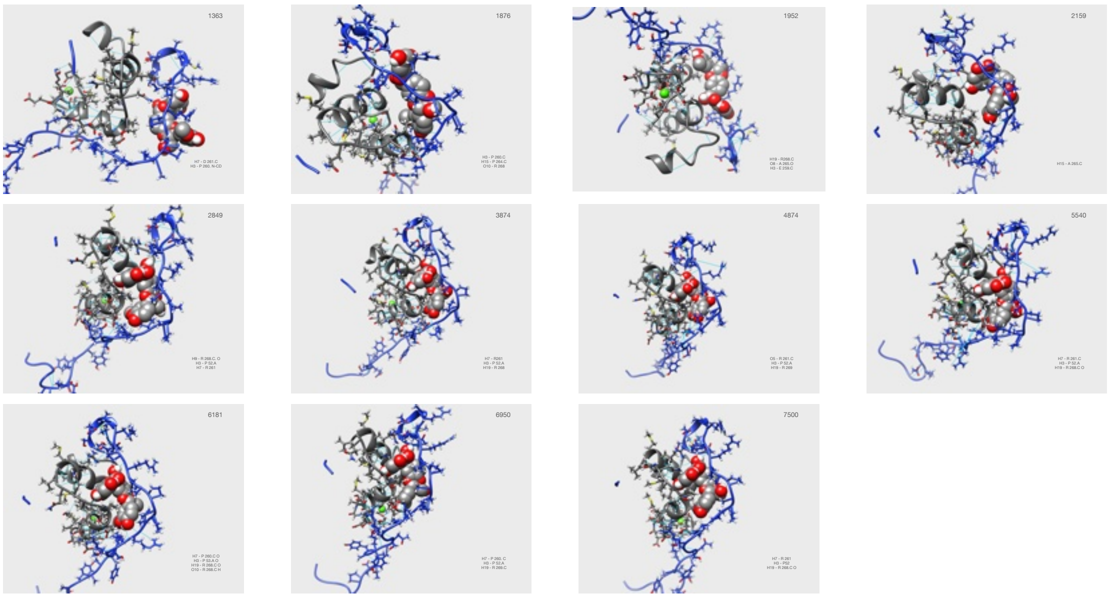

Silybin A

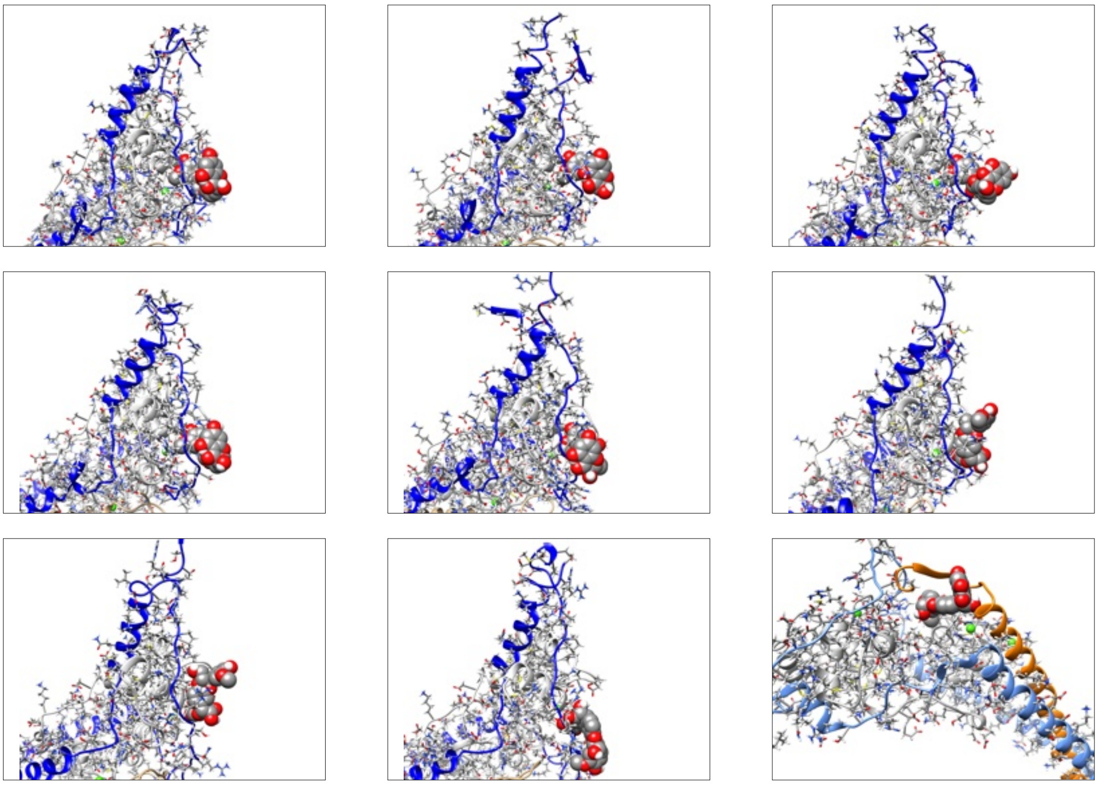

### EGCG

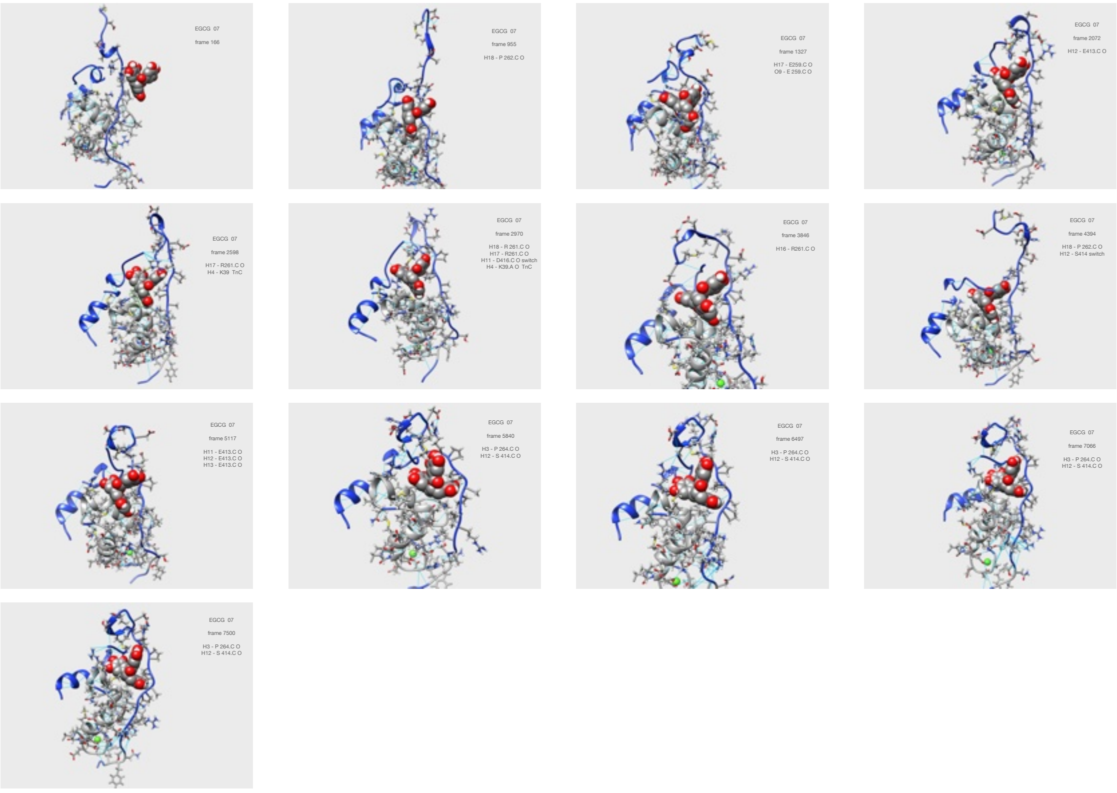

### Resveratrol

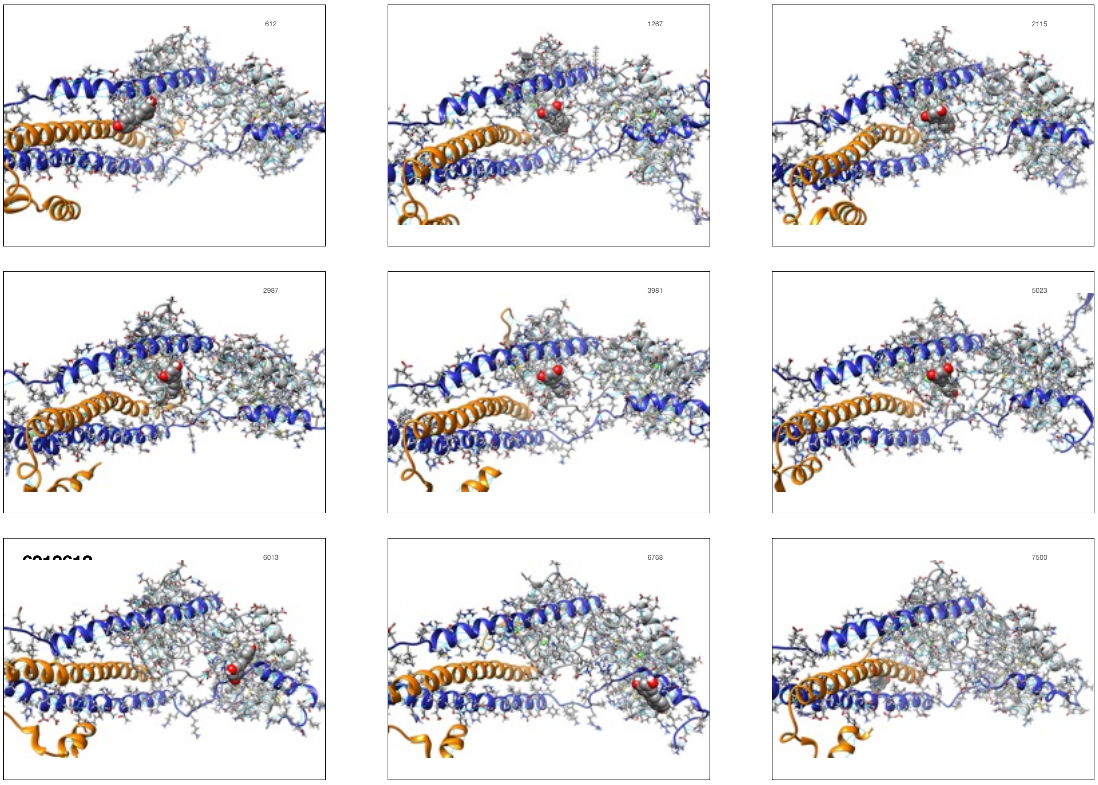
